# High Throughput Evolution of Near Infrared Serotonin Nanosensors

**DOI:** 10.1101/673152

**Authors:** Sanghwa Jeong, Darwin Yang, Abraham G. Beyene, Anneliese M.M. Gest, Markita P. Landry

## Abstract

Release and reuptake of neuromodulator serotonin, 5-HT, is central to mood regulation and neuropsychiatric disorders, whereby imaging serotonin is of fundamental importance to study the brain’s serotonin signaling system. We introduce a reversible near-infrared nanosensor for serotonin (nIRHT), for which synthetic molecular recognition toward serotonin is systematically evolved from ssDNA-carbon nanotube constructs generated from large libraries of 6.9 × 10^10^ unique ssDNA sequences. nIRHT produces a ∼200% fluorescence enhancement upon exposure to serotonin with a K_d_ = 6.3 µM affinity. nIRHT shows selective responsivity towards serotonin over serotonin analogs, metabolites, and receptor-targeting drugs, and a 5-fold increased affinity for serotonin over dopamine. Further, nIRHT can be introduced into the brain extracellular space in acute slice, and can be used to image exogenous serotonin reversibly. Our results suggest evolution of nanosensors could be generically implemented to rapidly develop other neuromodulator probes, and that these probes can image neuromodulator dynamics at spatiotemporal scales compatible with endogenous neuromodulation.

---

Serotonin, or 5-hydroxytryptamine (5-HT) is a monoamine neuromodulator that plays diverse roles in the central nervous system and is critically implicated in the etiology and treatment of mood and psychiatric disorders^1^. 5-HT also plays critical roles in modulating memory and learning^2, 3^, and in the regulation of mood^4^, sleep^5^, and appetite^5^. As such, aberrations in 5-HT signaling are thought to underlie multiple mental health disorders including major depressive disorder^6^, bipolar disorder^7^, autism^8^, schizophrenia^9^, anxiety^10^, and addiction^11^. Classical neurotransmission occurs through rapid activation of ligand-gated ion channels in which neurotransmitter signaling is confined at the synaptic cleft and thus to singular synaptic partners^12^. However, 5-HT acts primarily as a neuromodulator and may undergo signaling through extrasynaptic diffusion, enabling few neurons to affect broader networks of neuronal activity through 5-HT volume transmission. As such, there is great interest in monitoring the spatial component of 5-HT modulation in addition to its temporal dynamics. Current methods for measuring 5-HT include tools adapted from analytical chemistry to measure 5-HT in the brain extracellular space with varying levels of selectivity and spatiotemporal resolution: fast-scan cyclic voltammetry leverages redox chemistry to enable rapid temporal measurement of 5-HT by inserting a carbon electrode into brain tissue^13^. Microdialysis is an analytical chemistry method in which fluid samples from brain tissue are subject to downstream analysis with chromatography^14^. Recently, a field-effect transistor sensor modified with a 5-HT aptamer was reported to exhibit a highly sensitive and selective electrochemical response to 5-HT in physiological conditions^15^. However, previous methods suffer from limited spatial resolution, and microdialysis sampling occurs on the order of minutes, which precludes assaying fast 5-HT modulatory transients in the brain. While few optical probes for 5-HT of varying selectivity have been developed, to-date, none has been successfully implemented for monitoring 5-HT modulation on spatiotemporal scales relevant to monitoring endogenous 5-HT kinetics^16-21^. This lack of tools to probe 5-HT dynamics on commensurate spatiotemporal scales limits our ability to understand the role of 5-HT in both health and disease.

Herein, we report a reversible near-infrared optical probe for 5-HT that reports physiological 5-HT concentrations on relevant spatiotemporal scales, and is compatible with pharmacological tests. The probe responds with *ΔF/F*_*0*_ of up to 194% in the near-infrared fluorescence emission window of 1000-1300 nm, and is constructed from semiconducting single-walled carbon nanotubes (SWNT), which have previously shown utility for non-invasive through-skull imaging in rodents^22^. Molecular recognition and selectivity for 5-HT over its metabolic precursors, 5-HT metabolites, and 5-HT receptor drugs is conferred through surface adsorption of single-stranded polynucleotides to the SWNT surface. This near-infrared 5-HT probe (nIRHT) is developed through a nanosensor evolution technology introduced herein, in which large libraries of over 10^10^ polynucleotide-SWNT conjugates are evolved for 5-HT selectivity through iterative enzymatic amplification of 5-HT selective polymers and deep-sequencing to identify unique polynucleotide sequences that confer 5-HT selectivity when pinned to SWNT surfaces.

First, we describe a platform for evolution of molecular selectivity for neuromodulator 5-HT, in which systematic evolution of ligands by exponential enrichment is implemented on carbon nanotube surfaces, a process we’ve termed SELEC. While we’ve implemented SELEC to develop an optical probe for 5-HT herein, the platform is fundamentally generic for generating other optical probes of neurological relevance. We show that 6 rounds of polynucleotide-SWNT evolution enable generation of multiple candidate optical sensors for 5-HT with *ΔF/F*_*0*_ responses up to 194%. The best-responding construct from the evolution cohort, nIRHT, is next characterized as a 5-HT optical probe. nIRHT is shown to bind 5-HT with a K_d_ of 6.3 µM, is shown to be reversible, and exhibits unaltered performance in artificial cerebrospinal fluid (aCSF). Of importance to understanding pharmacology in the context of 5-HT signaling, we show that nIRHT does not respond to 5-HT metabolites of 5-hydroxyindoleacetic acid (HIAA), 5-hydroxytryptophan (HTP), and 5-methoxytryptophan (MTP) and 5-HT receptor-targeting drugs, such as fluoxetine, 3,4-methylenedioxymethamphetamine (MDMA), 25I-NMOMe, and quetiapine. Lastly, we demonstrate that nIRHT can be introduced into the brain extracellular space in acute striatal brain slices, and can be implemented to image exogenous 5-HT. Our results suggest nIRHT can serve as a probe to image 5-HT dynamics at spatiotemporal scales of relevance to endogenous 5-HT neurocommunication.

## RESULTS

### SELEC enables high-throughput screening of ssDNA-SWNT constructs for 5-HT neuromodulator sensitivity

Prior work has demonstrated that semiconducting near-infrared fluorescent SWNT, when noncovalently functionalized by amphiphilic polymers such as ssDNA, can generate synthetic molecular recognition elements on the SWNT surface. The resulting ssDNA-SWNT moiety can then selectively bind a molecular analyte for its optical detection^23-26^. The process of generating ssDNA-SWNT nanosensor candidates, and their screening against target analytes, has since revealed optical nanosensors for neurotransmitters including a (GT)_6_-SWNT nanosensor for dopamine^25^ which was subsequently used to image dopamine dynamics in the brain striatum and capture the influence of dopamine receptor drugs on dopamine dynamics at the level of individual synapses^27^. Despite the nascent utility of SWNT-based nanosensor technology, the nanosensor screening process involves a heuristic approach in which each nanosensor candidate must be synthesized individually before screening, limiting candidate nanosensor libraries to a few dozen, thus restricting the throughput of nanosensor discovery. Herein, we develop and validate an evolution-based approach for producing ssDNA-SWNT nanosensors in which upwards of 6.9 × 10^10^ unique ssDNA sequences are assessed for their ability to bind to a molecular analyte and produce an optical response. This strategy is inspired by SELEX protocols for aptamer generation^28^, and entails systematic evolution of ssDNA ligands by exponential enrichment following adsorption to carbon nanotubes (SELEC), which we implement to identify ssDNA-SWNT constructs that optically respond to neurotransmitter 5-HT. We designed our initial ssDNA library to be comprised of 6.9 × 10^10^ random 18-mer ssDNA sequences flanked by two (C)_6_-mers and two 18-mer primer regions for PCR amplification (Figure 1a). The 6-mer (C)_6_ sequence was introduced because this sequence was previously shown to have a high affinity for the SWNT surface^29^, and could thus serve as a SWNT-binding spacer sequence to separate the random 18-mer sequence from the PCR primer sequences. This ssDNA library was used to suspend SWNT using a previously described sonication protocol^25^, to create a SWNT suspension with 6.9 × 10^10^ unique ssDNA sequences for both an experimental and control ssDNA-SWNT sample: our experimental sample was generated in PBS solution by sonicating 10 µg SWNT with 2.8 mg ssDNA and 100 nmol of 5-HT. Our control sample was generated similarly but without 5-HT. The relative excess of ssDNA over SWNT was implemented to encourage competitive complexation of ssDNA to SWNT surfaces. Furthermore, the control sample served the role of identifying ssDNA sequences that are only strong SWNT surface binders, or subject to PCR biases^30^ during library preparation. In this manner, both experimental and control libraries could be generated for competitive binding of ssDNAs, followed by removal of unbound ssDNA sequences, and next desorption and deep sequencing of high-affinity ssDNA binders with an Illumina HiSeq 4000 (Figure 1a). For both the experimental and control libraries, we found that a total of six consecutive rounds of competitive ssDNA SWNT binding, desorption, recovery, and amplification of high-affinity ssDNA binders was sufficient to generate several promising 5-HT candidate nanosensors.

**Figure 1.**
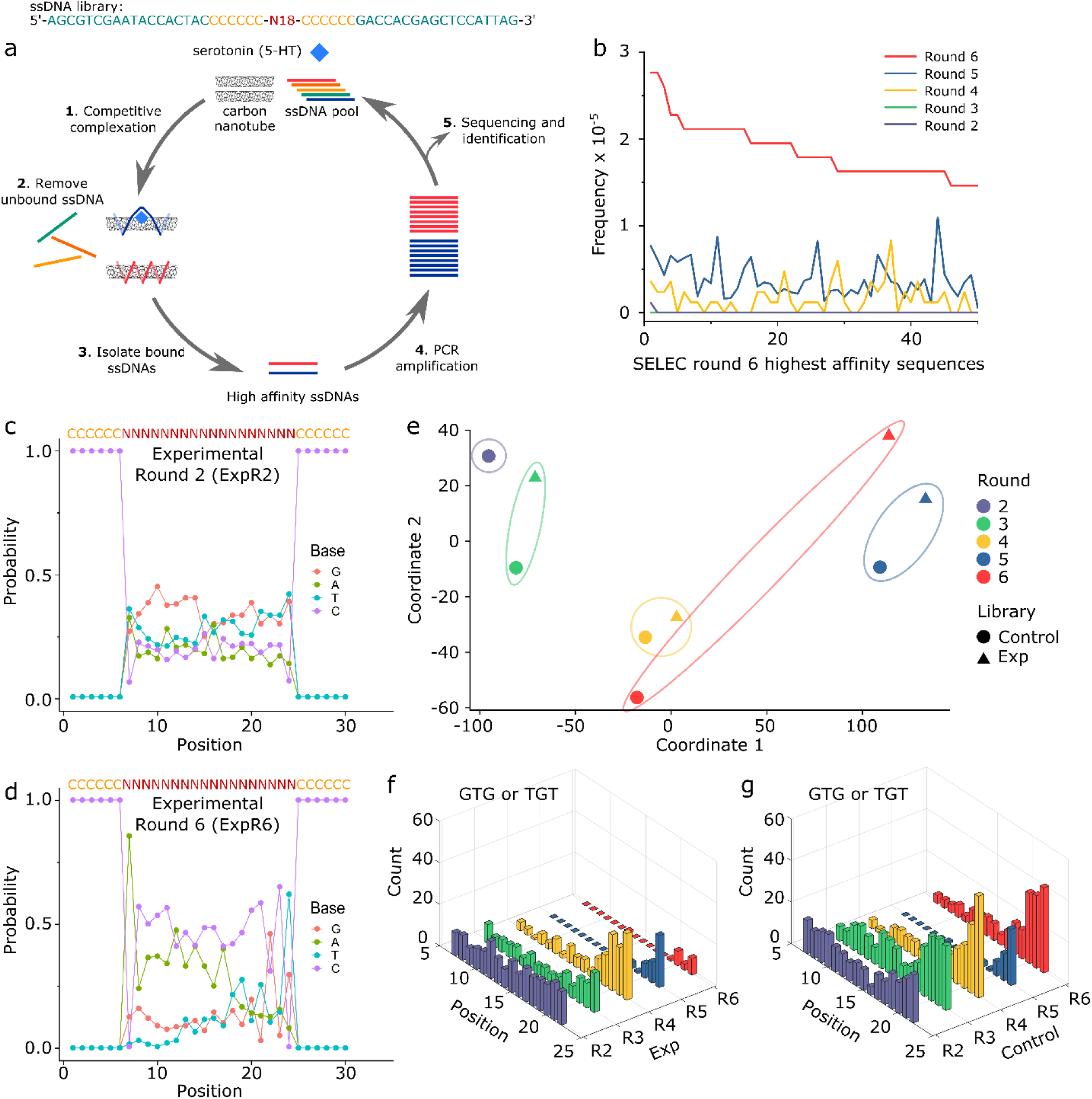
SELEC evolves ssDNA-SWNT nanosensors for 5-HT molecular recognition. (a) Schematic illustration of the SELEC evolution process. SWNTs are mixed with a >10-fold mass excess of ssDNA with 100 µM 5-HT (experimental library), or alone (control library). (1) The mixture is sonicated to generate the ssDNA-SWNT complex of either 5-HT-bound ssDNA-SWNT (blue) or ssDNA-SWNT (red). (2) Unbound ssDNAs are removed and (3) bound ssDNAs are isolated from SWNT by thermal desorption. (4) Isolated ssDNAs are amplified by PCR and the ssDNA pool is prepared for the next selection round and (5) characterized by high-throughput sequencing. (b) Sequence frequency of the top 50 unique sequences in the 6^th^ round of the experimental library, compared to their sequence prevalence in round 2, 3, 4, and 5. (c,d) Probability for each nucleotide at each position in the experimental library calculated from the top 200 sequences from (c) round 2 and (d) round 6. (e) Multidimensional scaling (MDS) plot of palindromic trimer frequencies of the top 200 sequences in the experimental and control SELEC libraries at each round. Data is plotted on the first and second coordinates of MDS analysis. Contour lines are included for better visualization of mutual divergence between experimental and control SELEC libraries for the same round. Divergence at round 2 may be too small to be distinguished. (f,g) Palindromic trimer incidences of (GTG)/(TGT) at each nucleotide position for (f) experimental and (g) control libraries.

The 50 most numerous thus the putatively highest-affinity sequences in the 6^th^ round of the experimental group were identified, and their corresponding frequencies in the total rounds were plotted in Figure 1b (see Table S1 and S2 for sequences of the top 10 binders). We observe a clear enrichment in the 50 highest-affinity sequences following the 6^th^ round of SELEC, in which each sequence has a mean frequency value of 19 × 10^-6^. Conversely, those sequences have mean frequency values of 1.6 × 10^-6^ and 3.8 × 10^-6^, respectively in round 4 and 5, and negligible or zero frequency preceding round 4. Next, the identity of each nucleotide base was evaluated with respect to position within the top 200 ranked sequences in each library. The round 2 experimental library showed a near random distribution of nucleotides at each position, with sequence identities near the ∼25% expected value at each position (Figure 1c). Following 6 rounds of SELEC, base identity preference trends emerge in the experimental (Figure 1d) and control (Figure S1) libraries. For instance, nucleotide A was preferred with a frequency of 86% and 65% in the experimental and control libraries at position 7, the first nucleotide following the 5’ (C)_6_ spacer. The experimental library round 6 sequences showed 65% C and 62% T identities at position 23 and 24, respectively, indicating a preference towards a 3’ (CT) terminus within the random region of the sequence. This preference is not observed in control library round 6 which instead showed enrichment of a (GNG) motif (Figure S1). Libraries were analyzed further by counting the incidences of nucleotide trimers with respect to position in the top 200 ranked sequences of each library. This analysis produced a trimer frequency table per nucleotide position for each control and experimental libraries. Classical multidimensional scaling (MDS) was performed on these frequency tables to evaluate the statistical difference between libraries. MDS reduces the dimensionality of library frequency tables into a two-dimensional (2D) projection, preserving the distances between trimer counts across libraries. A 2D MDS plot on 1^st^ and 2^nd^ principal coordinates showed that the experimental and control libraries at round 2 could not be distinguished, but the divergence between the experimental and control libraries became considerable by round 6 (Figure 1e). Round 6 divergence was driven in part by differences in abundance of repeating patterns such as (AC) and (GT) repeats. These differences were quantified by plotting counts of palindromic trimers such as (ACA) and (CAC), and (GTG) and (TGT) in the top 200 ranking sequences with respect to position (Figure S2, 1f, g). The repeating (AC) motif was preferred in both experimental and control SELEC groups and predominately located near the 5’ end. Interestingly, the repeating (GT) motif showed the largest contrast of motif population between experimental and control groups, which was selectively enriched at the 3’ end of the control group and not in the experimental group. We also performed direct principal component analysis (PCA) of the top 200 ranked sequences in each SELEC library. The 1^st^ and 2^nd^ principal component axes do not distinguish between the experimental and control libraries in the first four rounds, however, divergence is observed in round 5 and distinct clustering of the experimental from control libraries can be observed by round 6 (Figure S3). Together, these results suggest SELEC can evolve to select ssDNA sequences with high affinity for SWNT both in the presence and absence of an analyte, and that the sequences evolved in each are nearly mutually exclusive, with only one common sequence in the top 1000 sequences of both libraries at round 6. We therefore hypothesize that the experimental library will evolve ssDNA sequences that are selective for 5-HT when bound to SWNT, whereas the control library will evolve ssDNA sequences that exhibit highest affinity for the SWNT surface.

### In vitro characterization of nIRHT optical nanosensor identified from SELEC evolution

We next analyzed the sensitivity of SELEC evolved sequences to 5-HT when adsorbed to the surface of SWNT. We denote sequences in experimental group as E**N**#**M** (**M**th place sequence of round **N** in the order of frequency), and sequences from the control group as C**N**#**M**. ssDNA-SWNT nanosensor candidates were synthesized with the C_6_ anchor domains, but without the primers used for PCR-based amplification. Omission of primer regions from testing ssDNA-SWNT fluorescence responses was implemented to increase the mass density of the evolved N-18 library on the SWNT surface. This omission was performed following confirmation that both **C_6_-N18-C_6_** and **primer-C_6_-N18-C_6-_-primer** SELEC-evolved sequences from the experimental cohort showed enhanced sensitivity towards 5-HT, albeit with a reduced sensitivity for the latter, as predicted (Figure S4).

We prepared **C_6_-N18-C_6_** ssDNA-SWNT constructs from the top 10 most abundant sequences from the experimental and control groups for SELEC rounds 3, 4, 5, and 6 and tested the *ΔF/F*_*0*_ fluorescence response of each ssDNA-SWNT construct to 100 µM 5-HT in phosphate buffered saline (PBS) (Figure S5). *ΔF/F*_*0*_ is calculated as (F – F_0_)/F_0_ based on F_0_ of baseline fluorescence intensity before analyte addition and F of the fluorescence intensity 10 seconds after analyte addition for the (8,6) SWNT chirality (∼1195 nm)). As shown in Figure 2a, we find there is a noticeable increase in ssDNA-SWNT sensitivity for 5-HT as a function of SELEC round, relative to the baseline fluorescence modulation of *ΔF/F*_*0*_ = 0.73 for the ‘un-evolved’ ssDNA-SWNT sample prepared with the starting ssDNA library. This enhanced sensitivity towards 5-HT is most evident for the experimental library in which highly 5-HT sensitive constructs (*ΔF/F*_*0*_ > 1.2) are only found in the experimental SELEC group and not in the control group (Figure 2b). We identified the most 5-HT sensitive construct from both libraries and all SELEC rounds as E6#9 from the experimental library, which exhibited a *ΔF/F*_*0*_ of 1.94 ± 0.06 upon exposure to 100 µM 5-HT and could thus be used as a near-infrared 5-HT nanosensor (nIRHT). Multiple peaks in the fluorescence and absorption spectra of the nIRHT nanosensor showed the presence of different (n,m) SWNT chiralities in ensemble mixture^31^ (Figure 2c and Figure S6). We observe the turn-on response of nIRHT to 5-HT to be nearly instantaneous followed by a low fluorescence decay with τ_1/2_ > 100 s, which we confirmed is not a result of 5-HT structural or chemical change (Figure S7 and S8). For further characterization and use of nIRHT as a probe for 5-HT imaging, we focused on the optical response of (8,6) SWNT chirality corresponding to the emission peak at ∼1195 nm.

**Figure 2.**
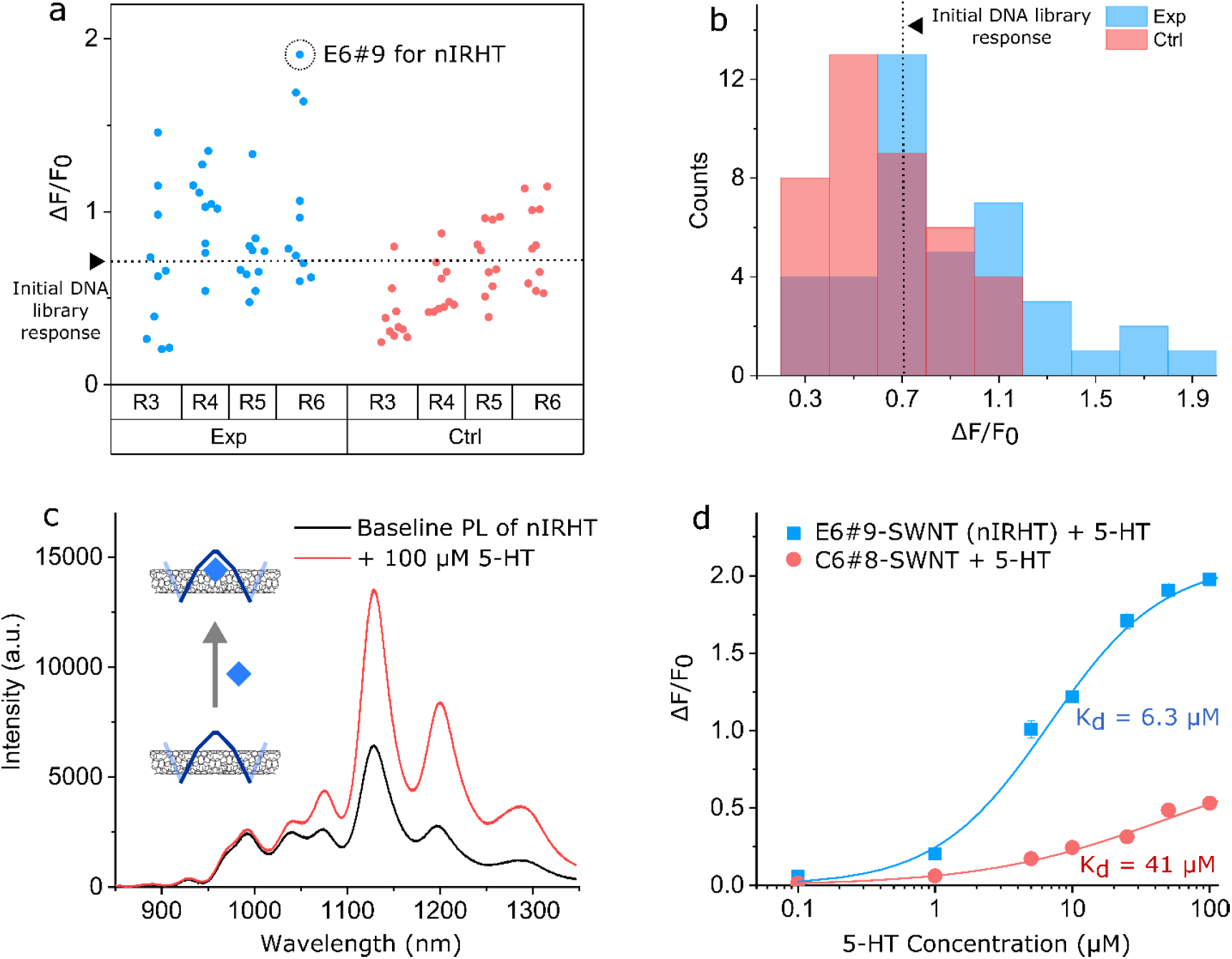
Evolution of ssDNA-SWNT demonstrates increased fluorescence sensitivity towards 5-HT. (a) *ΔF/F*_*0*_ upon addition of 100 µM 5-HT to top 10 ssDNA sequences from rounds 3, 4, 5, and 6 (R3, R4, R5, and R6) in experiment (Exp, blue circle) and control (Ctrl, red circle) SELEC groups (n = 3 trials). The most sensitive 5-HT nanosensor, E6#9, is indicated by a black dashed circle. *ΔF/F*_*0*_ from initial (round “zero”) ssDNA-SWNT library is represented by a dashed line. (b) Data from (a) represented as a histogram. (c) Fluorescence spectra of E6#9 ssDNA-SWNT before (black) and after (red) addition of 100 µM 5-HT. (d) *ΔF/F*_*0*_ 5-HT concentration dependence of E6#9-SWNT (blue) and C6#8-SWNT (red). Error bars denote standard deviation from n = 3 independent trials and may be too small to be distinguished in the graph. Experimental data are fitted with the Hill Equation (solid trace).

We next characterized nIRHT for use as a 5-HT brain imaging probe. We assessed the dynamic range of nIRHT to a 100 nM - 100 µM range of 5-HT concentrations, and show nIRHT sensitivity for 5-HT over a 100 nM to 50 µM dynamic range (Figure 2d), suitable for measuring endogenous 5-HT dynamics which are predicted to fall in the broad ∼100 pM - ∼1 µM concentration range^32-36^. Fitting nIRHT responsivity towards 5-HT to a Hill equation shows a micromolar dissociation constant (K_d_) of 6.3 µM for 5-HT (Figure 2d), a K_d_ previously identified as optimal for imaging neuromodulation to balance nanosensor reversibility for capturing fast transients^37^. We further confirmed that the ssDNA sequence for nIRHT has selective affinity for 5-HT through a solvatochromic shift assay with sodium cholate (Figure S9), and also confirmed that nIRHT shows a stable and reproducible fluorescence response to 5-HT for over a week post-synthesis (Figure S10). We next compared the sensitivity of nIRHT to an ssDNA-SWNT construct from the control SELEC group, C6#8, chosen for its strong affinity to the SWNT surface (Figure S9). When bound to SWNT, C6#8 showed reduced affinity (K_d_ = 41 µM) and a suppressed optical response (*ΔF/F*_*0*_ = 0.53 ± 0.01 at 100 µM 5-HT) compared to nIRHT (Figure 2d). These results suggest SELEC can be implemented to evolve ssDNA-SWNT selectivity for an analyte such as 5-HT, and that nIRHT can serve as an optical reporter for 5-HT imaging.

### Characterization of nIRHT as a nanosensor for 5-HT imaging in brain tissue

We next explored the utility of nIRHT for use as an optical probe to image extracellular 5-HT signaling in brain tissue. The response of nIRHT for 5-HT was assessed in aCSF, a common medium for imaging experiments in brain slice preparations. In aCSF, nIRHT demonstrated marginally superior performance than in PBS, with *ΔF/F*_*0*_ = 2.45 ± 0.07 upon exposure to 100 µM 5-HT and K_d_ = 10 µM (Figure 3a). We next examined the selectivity of nIRHT for 5-HT over other neurotransmitters. nIRHT nanosensor exhibited negligible *ΔF/F*_*0*_ response upon exposure to 100 µM acetylcholine, γ-aminobutyric acid, glutamate, tyrosine, and octopamine (Figure 3b). Dopamine, a catecholamine neurotransmitter, induced a moderate response of *ΔF/F*_*0*_ = 1.40 ± 0.03 upon addition of 100 µM dopamine, but nIRHT exhibited a 5-fold higher affinity for 5-HT over dopamine (K_d_ = 6.3 µM for 5-HT and K_d_ = 33 µM for dopamine) (Figure S11). We next measured nIRHT selectivity against 5-HT metabolites HTP, HIAA, and MTP, considering their high structural similarities to 5-HT. Addition of 100 µM HTP, HIAA, and MTP to nIRHT induced negligible fluorescence modulation of *ΔF/F*_*0*_ = 0.02 ± 0.02, 0.17 ± 0.10, and −0.14 ± 0.03, respectively. Lastly, given the relevance of 5-HT receptor drugs on the study of 5-HT modulation and pharmacology, we assessed selectivity of nIRHT against non-selective agonists fluoxetine and MDMA, 5-HT_2_ agonist 25I-NMOMe, and 5-HT_1A_ agonist quetiapine. Exposure of nIRHT to 100 µM fluoxetine, MDMA, 25I-NMOMe, and quetiapine induced negligible fluorescence modulation, and we additionally confirmed that 5-HT could be detected without attenuation even if nIRHT is pre-incubated with, and remains in the presence of, 1 µM of each of these drugs (Figure 3c).

**Figure 3.**
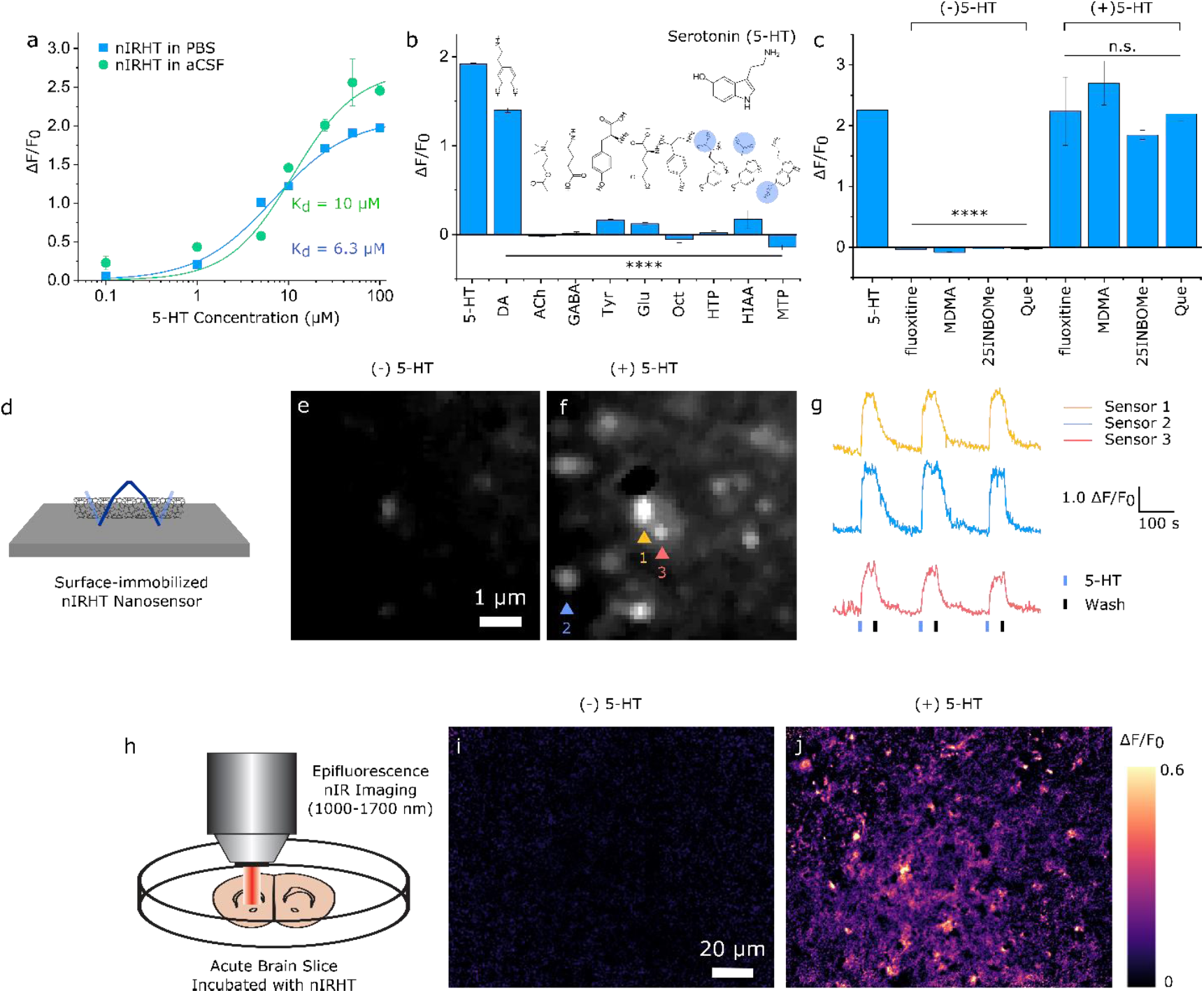
Validation and use of nIRHT 5-HT nanosensors in neurologically-relevant conditions. (a) 5-HT concentration dependent *ΔF/F*_*0*_ of nIRHT nanosensor in PBS (blue dot) and aCSF (green circle) fit with the Hill Equation (solid trace). (b) *ΔF/F*_*0*_ of nIRHT nanosensor upon exposure to 100 µM neurotransmitters and 5-HT metabolites (dopamine (DA), acetylcholine (ACh), γ-aminobutylic acid (GABA), tyrosine (Tyr), glutamate (Glu), octopamine (Oct), 5-hydroxy-L-tryptophan (HTP), 5-hydroxyindoleacetic acid (HIAA), 5-methoxytryptamine (MTP)). Error bars are standard deviation from n = 3 independent trials. (c) *ΔF/F*_*0*_ of nIRHT nanosensor upon exposure to 100 µM 5-HT with and without 1 µM 5-HT receptor drugs fluoxetine (non-selective agonist), 3,4-methylenedioxymethamphetamine (MDMA) (non-selective agonist), 25I-NMOMe (5-HT_2_ agonist), quetiapine (Que) (5-HT_1A_ agonist). Error bars are standard deviation from n = 3 independent trials. **** p < 0.0001 and n.s. means non-significant differences in one-way analysis of variance (ANOVA). (d) Reversibility of immobilized nIRHT nanosensors on glass substrate upon exposure to 100 µM 5-HT. nIR fluorescence images of the same field of view (e) before and (f) after addition of 100 µM 5-HT. (g) ΔF/F_0_ of immobilized nanosensors upon exposure to 100 µM 5-HT (blue bar) and rinsed with PBS (black bar) in a flow chamber. (h) near-infrared imaging of dorsomedial striatum from mouse acute brain slice after nIRHT nanosensor loading into brain tissue. (i,j) *ΔF/F*_*0*_ images of the same field of view of an acute striatal brain slice (i) before and (j) after exogenous addition of 100 µM 5-HT.

Towards endogenous 5-HT imaging, we next investigated the reversibility of the nIRHT nanosensor by immobilizing nIRHT in a microfluidic slide, and imaging individual nIRHT nanosensors upon successive infusions of 100 µM 5-HT. nIRHT nanosensors were immobilized onto a (3-aminopropyltriethoxysilane) (APTES)-treated glass slide, and placed in a flow chamber with a syringe pump. The nanosensor chamber was exposed to three cycles of 100 µM 5-HT in PBS subsequently rinsed with PBS only, where the near-infrared fluorescence response was imaged on an inverted microscope as previously described^27^ (Figure 3d-f, and Supplementary Movie 1). We observe nIRHT fluorescence increase and decrease as predicted by the influx and washing of 5-HT, respectively. Specifically, the time-dependent near-infrared fluorescence of three individual nanosensors was measured across three cycles and were found to be *ΔF/F*_*0*_ = 1.55 ± 0.01, 1.71 ± 0.08, and 1.10 ± 0.14 (mean ± SD for three cycles), which indicates that nIRHT nanosensors are reversible (Figure 3g). We suggest the variation in *ΔF/F*_*0*_ among nanosensors is a result of the chirality-dependent variability in SWNT response as observed in bulk measurements. We further demonstrated that surface-immobilized SWNT, without the ssDNA coating, do not respond to 5-HT, confirming the molecular recognition role of the ssDNA polymer when adsorbed to the SWNT surface (Figure S12). Importantly, prior studies of ssDNA-SWNT nanosensors have confirmed ms-scale nanosensor reversibility kinetics^25, 27^, the absence of photobleaching^38^, and *in vivo* compatibility^39^, suggesting nIRHT could serve as an optical probe for long-term endogenous 5-HT imaging in the brain.

Lastly, we tested nIRHT nanosensor compatibility for 5-HT imaging in acute brain slices. Coronal mouse brain slices of 200 µm thickness were incubated with a 2 µg/mL nIRHT suspension for 15 minutes and subsequently rinsed with aCSF to remove excess nIRHT. This labeling procedure was previously found to label acute slices with nanosensor localization in the brain extracellular space^27^. We next placed the labeled slice in an imaging chamber in aCSF for imaging on a custom-built nIR fluorescence upright epifluorescence microscope (Figure 3h). Time-dependent fluorescence images were recorded to confirm stable near-infrared fluorescence of nanosensors when no external stimuli were present (Supplementary Movie 2). We then investigated the ability of nIRHT nanosensors to respond to 5-HT in slice. Following acquisition of one minute fluorescence baseline, bath application of 100 µM 5-HT resulted in a nIRHT fluorescence increase of *ΔF/F*_*0*_ = 0.35 ± 0.26; mean ± SD (Figure 3i, j, and S13 and Supplementary Movie 3). Rinsing of the slice imaging chamber with aCSF confirmed that nIRHT fluorescence when in slice returns to baseline in the absence of 5-HT.

## DISCUSSION

High spatiotemporal imaging of neuromodulatory signaling is essential to understand the dynamics of neuromodulator release and uptake in neural circuits and to elucidate the function of chemical neuromodulation in animal behavior. Synthetic probes are an attractive approach to imaging neuromodulators such as 5-HT owing to their photostability, compatibility with pharmacology, and feasibility for use in genetically-intractable organisms. Herein, we develop an approach to discover near infrared optical probes through evolution of ssDNA-SWNT selectivity towards 5-HT. ssDNA-SWNT constructs whose polymer corona phase recognizes 5-HT were evolved from an initial 18-mer nucleotide library of > 10^10^ unique sequences by inducing competitive complexation between ssDNAs and SWNTs in the presence of 5-HT, through a process we termed SELEC. Prior studies have shown that analytes that induce selective fluorescence modulation of SWNTs also stabilize the ssDNA corona^25^, thus we reason that evolved 5-HT recognizing sequences may preferentially populate the evolved ssDNA pool via stabilization of the 5-HT-ssDNA-SWNT constructs. Conversely, the control SELEC library evolved in the absence of 5-HT should contain ssDNA sequences with high affinity for the SWNT surface but not specifically for 5-HT. Indeed, we find that pE6#9-SWNT (the 5-HT responsive nIRHT nanosensor with PCR primer regions) is stabilized by the presence of 5-HT substantially more so than pR0-SWNT (the initial un-evolved ssDNA library) (Figure S9). This result further indicates the ability of SELEC to evolve ssDNA-SWNT nanosensors for selective recognition of analytes such as 5-HT. Taken together, these data demonstrate that SELEC can be implemented to rapidly screen large libraries of ssDNA-SWNT constructs to evolve selectivity for small biomolecule analytes, circumventing low-throughput screening of such constructs previously used for SWNT-based nanosensor development. It is feasible that SELEC could be implemented for other ssDNA-SWNT technologies such as SWNT-based chirality separation^31, 40, 41^, which is strongly dependent on SWNT surface adsorption of specific ssDNA sequences. Furthermore, because SELEC is fundamentally agnostic to the biomolecule analyte(s), SELEC may be implemented to develop near-infrared optical probes for other small molecule targets of interest.

By implementing SELEC to evolve molecular recognition for neuromodulator 5-HT, we identified a nanosensor, nIRHT, with *ΔF/F*_*0*_ of 1.94 ± 0.06 in PBS, and *ΔF/F*_*0*_ of up to 2.45 ± 0.07 in aCSF. Furthermore, we demonstrated that nIRHT provides a selective optical response for 5-HT against 5-HT metabolites, other neuromodulators, and pharmacology of relevance to 5-HT neuromodulation. While neurotransmitter dopamine also induced a moderate *ΔF/F*_*0*_ = 1.36 fluorescence increase of nIRHT (Figure 3b), nIRHT exhibits a 5-fold greater affinity for 5-HT over dopamine with K_d,5-HT_ = 6.3 µM and K_d,dopamine_ = 33 µM (Figure S11). The SELEC process therefore shows promise for evolution of selective optical probes, which may also benefit from the evolution of a positive control library to generate enrichment of ssDNA-SWNT constructs that are responsive to cross-reactive molecules. This additional ‘positive’ selection could help identify sequences with domains uniquely responsive to interfering molecules and eliminate those motifs from positive hits for the target ligand. It is noteworthy that the synthetically evolved nIRHT nanosensor shows negligible cross-responsivity with 5-HT precursors and metabolites, and 5-HT receptor agonists and antagonists. Many optical probes for neurochemicals are based on proteins engineered from endogenous receptors and thus show cross-reactivity with receptor-targeting drugs. However, because the molecular recognition sites of nIRHT nanosensors is synthetic in nature, we demonstrate that nIRHT nanosensors do not show fluorescence modulation upon exposure to 5-HT receptor drugs fluoxetine, MDMA, 25I-NMOMe, and quetiapine (Figure 3c). Thus, nIRHT optical probes and others of its class could enable insights into synaptic-level effects of 5-HT receptor agonists and antagonists, to further our understanding of pharmacology of relevance to major depressive disorder^6^, bipolar disorder^7^, autism^8^, schizophrenia^9^, anxiety^10^, and addiction^11^ among others.

In comparison to the reported methodologies for 5-HT dynamic imaging such as FSCV^13^, microdialysis^14^, and paramagnetic 5-HT binding proteins for MRI^42^, nIRHT nanosensors are promising high spatiotemporal fluorescence imaging probes for studying 5-HT neuromodulation, which provide sub-micrometer spatial resolution, reversibility, photostability, and selectivity against 5-HT metabolites and receptor-targeting drugs. Furthermore, SWNT fluoresce in a unique near-infrared window (1000-1300 nm) which reduces photon scattering and enhances imaging penetration depths, enabling through-cranium^22^ and *in vivo* imaging^39^. This optical window could enable the through-cranium fluorescence imaging of cortical 5-HT neuromodulation without a cranial window, minimizing surgical perturbations for chronic 5-HT imaging in animal models with high biocompatibility^22, 43^. Furthermore, prior work has demonstrated that SWNT-based neuromodulator probes are tractable for use in non-model species^27^, further expanding the repertoire of neurological studies to be undertaken with nIRHT and other probes of its class.

## METHODS

### Materials

HiPCo single-walled carbon nanotubes (SWNT) were purchased from NanoIntegris (Batch #27-104). serotonin hydrochloride (hydroxytryptamine hydrochloride), dopamine hydrochloride, acetylcholine, γ-aminobutylic acid (GABA), tyrosine, glutamate, octopamine, 5-hydroxy-L-tryptophan (HTP), 5-hydroxyindoleacetic acid (HIAA), 5-methoxytryptamine (MTP), fluoxetine, 3,4-methylenedioxymethamphetamine (MDMA), 25-NMOMe, and quetiapine were purchased from Sigma Aldrich. All ssDNA sequences were purchased from integrated DNA technologies (IDT, USA).

### Optical characterization of SWNT

nIR fluorescence spectra were measured with a 20 X objective on an inverted Zeiss microscope (Axio Observer.D1) coupled to a spectrograph (SCT 320, Princeton Instruments) and liquid nitrogen cooled InGaAs linear array detector (PyLoN-IR, Princeton Instruments). A 721 nm laser (OptoEngine LLC) was used as the excitation light source for all spectroscopy and imaging studies. The absorption spectra were measured with a UV-3600 Plus absorption spectrophotometer (Shimadzu). Our custom near-infrared spectrometer and microscope have been described in detail in previous works^44, 45^.

### SELEC experimental protocol

The initial ssDNA random library was purchased from integrated DNA technologies (IDT, USA), which consisted of 18 random nucleotides flanked by two fixed 6-mer (C)_6_ and two 18-mer primer regions for PCR amplification: AGCGTCGAATACCACTAC-CCCCCC-N18-CCCCCC-GACCACGAGCTCCATTAG. For the first round, 100 nmol of the ssDNA library was dissolved in PBS buffer, and the mixture was denatured by heating at 95 °C for 5 min and subsequently cooled to room temperature. For the experimental SELEC group to evolve 5-HT molecular recognition, the ssDNA library was incubated with 100 µM 5-HT for 10 min in 0.99 mL PBS buffer. Next, 10 µg SWNT was added to the ssDNA and 5-HT mixture. A separate SELEC control group was prepared as described above but without addition of 5-HT. The resulting mixtures were bath-sonicated for 2 min and next probe tip-sonicated (Cole Parmer Ultrasonic Processor, 3 mm tip) for 10 min at 5 W power in an ice bath. After sonication, the black SWNT suspension was centrifuged for 60 min at 16,100 g to precipitate any un-suspended SWNT, and the supernatant containing the ssDNA-SWNT constructs solution was collected. The supernatant was spin-filtered using a 100 kDa molecular weight cutoff (MWCO) centrifugal filter (Amicon Ultra-0.5, Millipore) at 6,000 rpm for 5 min with DNase free water to remove unbound ssDNAs and 5-HT, and the remaining solution was collected. The spin filtration was repeated five times. Next, the purified ssDNA-SWNT suspension was heated at 95 °C for 1 h to detach surface-bound ssDNAs from SWNT. Following heating, black SWNT aggregates were observed, as expected when the ssDNA corona desorbs from the SWNT surface. The ssDNA solution with aggregated SWNT was centrifuged for 10 min at 16,100 g to further pellet undissolved SNWT, and the supernatant containing ssDNA was collected. The collected ssDNA library was amplified by PCR using a FAM-modified forward primer (FAM-AGCGTCGAATACCACTAC) and biotinylated backward primer (biotin-CTAATGGAGCTCGTGGTC), following a previously described protocol^28^. Standard solution conditions for PCR amplification were utilized as follows, to maximize dsDNA yield while reducing by-products after amplification: 2.5 units Hot Start Taq DNA polymerase (New England Biolabs (NEB)), 1X Hot Start Taq reaction buffer, 1 µM forward primer, 1 µM backward primer, 500 µM dNTP, ∼100 ng/mL ssDNA library template in a total 10 mL volume for each of the 100 µL volume 96 reaction wells. A negative control well was prepared without ssDNA library template. A preparative PCR reaction was performed with the following standard cycling conditions: initial denaturation for 150 s at 95 °C; N cycles of denaturation for 30 s at 95 °C and annealing for 30 s at 50 °C and extension for 30 s at 72 °C; final extension for 180 s at 72 °C. The cycle number N is determined before preparative PCR, which yielded maximal ssDNA and negligible PCR by-product formation (usually between 20 to 30 cycles). Next, 100 µL of the PCR products from each the experimental and the control SELEC libraries were separately collected and purified with a GeneJET PCR purification kit (Thermo Scientific) for preparation of high-throughput sequencing libraries. PCR products were confirmed by gel electrophoresis in a 4% agarose gel (low range ultra agarose, Bio-Rad laboratories) in 1X Tris-borate-EDTA (TBE) buffer (run for 18 min at 110 V). After electrophoresis, the gel was stained with SYBR gold and DNA bands were observed under UV light. To generate ssDNA from PCR products, 2.5 mL of streptavidin-coated beads (Streptavidin Sepharose High Performance, GE Healthcare) was placed on a sintered glass buchner funnel (pore size < 10 µm) and washed with 10 ml PBS buffer. The PCR product solution was incubated with the beads for 30 min and passed through the funnel. The incubation step was repeated three times to bind dsDNA onto the beads, and the beads were washed again with 10 ml of water. To elute the FAM-labeled ssDNAs, 8 ml of 0.2 M NaOH aqueous solution was added slowly to the beads, and the eluate was collected containing the FAM-labeled ssDNAs. The eluted solution was desalted with an NAP10 desalting column (GE Healthcare), and concentrated using a DNA speedvac dryer. The amount of ssDNA was quantified by absorbance at 260 nm, and usually 2-5 nmol of ssDNA was obtained and utilized for the next round of SELEC. For the control SELEC library evolved with only ssDNA and SWNT in the absence of 5-HT, the steps above were performed without 5-HT coincubation.

### High-throughput sequencing and analysis

Sequencing libraries were prepared by using two sequential PCR steps^46^ to add Illumina TruSeq universal adapter sequences and sequencing indices for multiplexed sequencing of round 2, 3, 4, 5, and 6 from experimental and control SELEC groups. The sequencing libraries was sequenced with an Illumina HiSeq 4000 at the Genomic Sequencing Laboratory in the University of California, Berkeley, which generated about 20M raw sequences from each round. We filtered out sequences that did not contain correct fixed regions (18-mer from 5’ primer + 6-mer cytosine block + 18-mer random region + 6-mer cytosine block + 3-mer from 3’ primer). A FASTAptamer toolkit was utilized to filter out and determine sequence frequencies^47^. In the manuscript, all experiments used ssDNA sequences comprising only the 18-mer random region flanked by 6-mer cytosine blocks at each end (without PCR primer regions), unless otherwise noted. For classical multidimensional scaling (MDS), the frequencies of trimers (e.g. ACA) at each position in the top 200 ranking sequences from the experimental or control SELEC libraries at each round were calculated and used as input parameters. Two dimensional MDS analysis was performed by using the cmdscale() function of the stats package in R.

### ssDNA-SWNT construct preparation for nanosensor characterization

To characterize nanosensor properties such as fluorescence sensitivity and selectivity to 5-HT, ssDNA-functionalized SWNT constructs were generated with the following protocol: 1 mg of HiPCo SWNT was added to 0.9 mL of PBS buffer, and the solution was mixed with 100 µL of 1 mM ssDNA. The resulting mixture was bath-sonicated for 2 min and tip-sonicated for 10 min at 5 W power in an ice bath. After sonication, the black ssDNA-SWNT suspension was centrifuged for 30 min at 16,100 g to precipitate non-dispersed SWNT, and the supernatant containing solubilized ssDNA-SWNT was collected. The supernatant was spin-filtered with 100 kDa MWCO centrifugal filters at 6000 rpm for 5 min with DNase free water to remove unbound ssDNAs, and the purified solution at the top of the filter was collected. This spin filtration to remove unbound ssDNA was repeated three times. The ssDNA-SWNT suspension was diluted with PBS buffer, and stored at 4 °C until use. The concentration of the ssDNA-SWNT suspension was calculated by measuring its absorbance at 632 nm with an extinction coefficient for SWNT^26^ of 0.036 (mg/L)^-1^ cm^-1^.

### Spectral solvatochromic shift of ssDNA-SWNT induced by sodium cholate (SC)

For SC-induced spectral shift assays without 5-HT, ssDNA-SWNT samples were prepared by creating a 100 µL PBS solution with 20 mg/L ssDNA-SWNT generated with the following ssDNAs: pR0, pE6#9, and pC6#8 ssDNAs containing the PCR primer regions (Table S3). These solutions were loaded onto a custom-built nIR fluorescence spectrometer and the time-dependent nIR fluorescence spectra were measured before and after addition of 100 µL of 0.5 wt% SC in PBS. The spectral blue-shift was observed for all the fluorescence peaks, and we quantified the time-dependent spectral shift of the (8,6) SWNT chirality peak (∼1195 nm). For the SC-induced spectral shift assay with 5-HT pre-incubation, a 100 µL PBS solution containing 20 mg/L ssDNA-SWNT of the above-mentioned sequences was incubated in 100 µM 5-HT for 1 minute. The fluorescence spectra of the ssDNA-SWNT constructs was subsequently measured every 2 seconds for a total of ∼6 minutes, with 0.5 wt% SC added to each sample at a timepoint marked by a solid black arrow (Figure S9).

### Screening the fluorescence response of ssDNA-SWNT constructs to 5-HT and other analytes

Fluorescence spectra of 198 µL 10 mg/L ssDNA-SWNT suspensions in PBS were measured before and 10 s after the addition of 2 µL of 10 mM 5-HT solution, for a final 5-HT concentration of 100 µM. We analyze the nIR fluorescence change of the (8,6) SWNT chirality peak (∼1195 nm) in this study. ΔF/F_0_ was calculated as *ΔF/F*_*0*_ = (F – F_0_)/F_0_ based on the baseline fluorescence intensity before analyte addition F_0_, and the fluorescence intensity 10 s after analyte addition for the (8,6) SWNT chirality (∼1195 nm), F. Most of the analyte solutions were prepared with dissolution in DI water as 10 mM concentration stocks, except tyrosine which was dissolved in DMSO. Serotonin hydrochloride and dopamine hydrochloride were used to prepare the analyte solutions to prevent oxidation of the neuromodulators in ambient conditions, and the hydrochloride salt did not affect the pH of resulting mixture because the PBS buffer capacity (∼10 mM) is significantly higher than the amount of hydrochloride in the neuromodulator stocks (0.1 mM). All *ΔF/F*_*0*_ measurements were performed in triplicate, and the mean and standard deviation of triplicate measurements were plotted. The assessment of 5-HT metabolites on nIRHT fluorescence was tested by preparing a 198 µL 10 mg/L nIRHT suspension in PBS and taking the nIR fluorescence spectra before and 10 s after addition of 2 µL of 10 mM 5-HT metabolites (HTP, HIAA, MTP). The assessment of 5-HT receptor drugs on nIRHT fluorescence was tested by preincubating a 198 µL of 10 mg/L nIRHT suspension in 1 µM drug (fluoxetine, MDMA, 25I-NMOMe, and quetiapine) in PBS and taking the nIR fluorescence spectra before and after addition of 2 µL of 10 mM 5-HT. All concentration-dependent *ΔF/F*_*0*_ response curves were fitted to the Hill equation^24^ to determine K_d_’s. To measure the optical ΔF/F_0_ response of nIRHT in brain-relevant environments, we prepared a 10 mg/L nIRHT stock solution in artificial cerebrospinal fluid (aCSF) composed of 119 mM NaCl, 26.2 mM NaHCO_3_, 2.5 mM KCl, 1 mM NaH_2_PO_4_, 1.3 mM MgCl_2_, 10 mM Glucose, 2 mM CaCl_2_ at pH 7.4. In a separate assay, 100 µL of 100 mg/L nIRHTs were incubated in 900 µL of stock human cerebrospinal fluid (CSF, purchased from Lee biosolution).

### NMR analysis of 5-HT after coincubation with nIRHT

For NMR analysis, PBS buffer was prepared in D_2_O (PBS-d). 396 µL of 10 mg/L nIRHT in PBS-d was mixed with 4 µL of 10 mM 5-HT in D_2_O. The mixture was excited with a 721 nm laser for 10 min on the same nIR fluorescence spectrometer used for nanosensor screening and fluorescence measurements. The solution was spin-filtered using a 100 kDa MWCO centrifugal filter at 6,000 rpm for 5 min with D_2_O, and the flow-through solution was collected. The ^1^H NMR spectra of the flow-through solution was measured with a 400 MHz Avance AVB-400 NMR (Bruker). The ^1^H NMR spectra of a 0.1 mM 5-HT solution without laser excitation in D_2_O was also measured as a control.

### Reversibility of nIRHT nanosensors immobilized in a microfluidic chamber

A glass coverslip (#1.5, thickness = 0.17 mm, HS159879H, Heathrow Scientific) was functionalized with (3-Aminopropyl) triethoxysilane (APTES, Sigma Aldrich) by soaking in 10 v/v% APTES in anhydrous ethanol for 5 min, following a previously reported protocol^27^. The coverslip was then rinsed with DI water and left to dry. The coverslip was then fixed onto an ibidi µ-Slide VI^0.5^ forming 6 microfluidic channels. Next, 100 μL of PBS was pipetted through a microfluidic channel. The channel was then filled with 50 μL of a 10 mg/L solution of nIRHT and left to incubate at room temperature for 5 min. The channel was rinsed using three successive washes of 50 μL PBS. The surface immobilized nIRHT in PBS were then imaged on an inverted light microscope with 721 nm excitation and a Ninox VIS-SWIR 640 camera (Raptor photonics). One end of the flow channel was connected to a syringe pump (Harvard Appartus) with Luer lock fittings. Prior to the start of image acquisition, the opposite flow reservoir was filled with PBS and the pump was set to refill mode at a volumetric flow rate of 100 μL/min. Once the liquid in the reservoir was depleted, 50 μL of 100 μM 5-HT in PBS was added. The process was repeated using alternating additions of 200 μL PBS and 50 μL of 5-HT solution.

To test the optical response of bare SWNT immobilized onto a glass substrate, an SDS-coated SWNT suspension was prepared: 1 mg of HiPCo SWNT was added to 1 mL of 2 wt% SDS in PBS buffer. The resulting mixture was bath-sonicated for 2 min and tip-sonicated for 30 min at 5 W power in ice bath. After sonication, the black suspension was centrifuged for 60 min at 16,100 g to precipitate undissolved SWNT, and the supernatant solution was collected. 50 µL of 5 mg/L SDS-SWNT was incubated in a microfluidic slide as described above, and the SDS coating was gradually removed by introducing a continuous flow of PBS buffer into the channel for 1 h at a flow rate of 50 µL/min. Following this surfactant stripping step, 50 μL of 10 μM 5-HT in PBS was added to the microfluidic channel.

### nIRHT nanosensor imaging in an acute brain slice

60 day old C57 Bl/6 strain mice were used for nIRHT brain slice imaging experiments. Mice were group housed after weaning at P21 and kept with nesting material on a 12:12 light cycle. All animal procedures were approved by the UC Berkeley Animal Care and Use Committee (ACUC). Acute brain slices were prepared using established protocols^48^. Briefly, mice were deeply anesthetized via intraperitoneal injection of ketamine/xylazine cocktail and transcardial perfusion was subsequently performed using ice-cold cutting buffer (119 mM NaCl, 26.2 mM NaHCO_3_, 2.5 mM KCl, 1mM NaH_2_PO_4_, 3.5 mM MgCl_2_, 10 mM glucose, 0 mM CaCl_2_), after which the brain was rapidly extracted. The cerebellum and other connective tissues were trimmed using a razor blade and the brain was mounted onto the cutting stage of a vibratome (Leica VT 1200S). Coronal slices (300 µm in thickness) including the dorsal striatum were prepared. Slices were incubated at 37°C for 60 minutes in oxygen saturated aCSF (119 mM NaCl, 26.2 mM NaHCO_3_, 2.5 mM KCl, 1mM NaH_2_PO4, 1.3 mM MgCl_2_, 10 mM glucose, 2 mM CaCl_2_) before use. Slices were then transferred to room temperature for 30 minutes before starting imaging experiments and were maintained at room temperature for the remainder of experimentation. For nanosensor labeling, slices were transferred into a small volume brain slice incubation chamber (Scientific Systems Design, Inc., AutoMate Scientific) and kept under oxygen saturated aCSF (total 5 mL volume). 100 µL of 100 mg/L nIRHT nanosensor was added to the 5 mL volume and the slice was incubated in this solution for 15 minutes. The slice was subsequently recovered and rinsed in oxygen saturated aCSF to wash off nIRHT that did not localize into the brain tissue. The rinsing step was performed by transferring the slice through 3 wells of aCSF in a 24 well plate (5 seconds in each well) followed by labeled slice transfer to the recording chamber with aCSF perfusion for a 15-minute equilibration period before starting the imaging experimentation. All imaging experiments were performed at 32 °C with continuous perfusion of oxygen-saturated aCSF to maintain brain slice viability. *Ex vivo* slice imaging was performed with a modified upright epifluorescensce microscope mounted onto a motorized stage as we previously described^27^. The brain slice labeled with nIRHT was illuminated with 785 nm light, and the fluorescence images were taken in a field of view within the brain dorsomedial striatum with an InGaAs array detector (Ninox 640, Raptor) with 100 ms exposure and a 60 X objective. To observe nIRHT response to 5-HT influx, the brain slice was immersed in 3 mL of oxygen saturated aCSF in the flow chamber and 0.3 mL of 1 mM 5-HT in aCSF was added into the system without further mixing. After the fluorescence reached a plateau, the 5-HT in the chamber solution was slowly depleted by continuous flow of oxygen saturated aCSF (washing). Time-dependent ΔF/F_0_ was calculated from multiple region of interests (ROIs) identified by a custom image analysis program (available for download: https://github.com/jtdbod/Nanosensor-Brain-Imaging).

## ACKNOWLEDGEMENTS

We acknowledge support of an NIH NIDA CEBRA award # R21DA044010 (to M.P.L.), a Burroughs Wellcome Fund Career Award at the Scientific Interface (CASI) (to M.P.L.), the Simons Foundation (to M.P.L.), a Stanley Fahn PDF Junior Faculty Grant with Award # PF-JFA-1760 (to M.P.L.), a Beckman Foundation Young Investigator Award (to M.P.L.), and a DARPA Young Investigator Award (to M.P.L.). M.P.L. is a Chan Zuckerberg Biohub investigator. This work used the Vincent J. Coates Genomics Sequencing Laboratory at UC Berkeley, supported by NIH S10 OD018174 Instrumentation Grant.

## AUTHOR CONTRIBUTIONS

S.H.J. and M.P.L. conceived the ideas of SELEC and designed experiments. S.H.J. and A.M.M.G. performed the iterative process of SELEC and characterized the nanosensor properties. D.Y. and S.H.J. analyzed the high throughput sequencing data. A.G.B. and S.H.J. carried out *ex vivo* acute brain slice experiments. S.H.J., D.Y., A.G.B., A.M.M.G., and M.P.L. discussed the experimental results and wrote the manuscript. All authors discussed the results and commented on the manuscript.

