## Supplemental Information for "High Throughput Evolution of Near Infrared Serotonin Nanosensors"

Jeong *et al.*

| Name | Sequence | Note |
|---|---|---|
| E3#1 | CCCCCCACCTGACACGATCCTATGCCCCCC | |
| E3#2 | CCCCCCGGCACAACGCTCGATGCTCCCCCC | |
| E3#3 | CCCCCCATTACAGCGGACAAGTGTCCCCCC | |
| E3#4 | CCCCCCTAAGGCCGATCCCACTATCCCCCC | |
| E3#5 | CCCCCCTGACTCCATAACAGTGTGCCCCCC | |
| E3#6 | CCCCCCGACACCCTGGACCCGTCGCCCCCC | |
| E3#7 | CCCCCCTGGCGTACAAACCGTCTGCCCCCC | |
| E3#8 | CCCCCCACACACTCTACTCTTCCACCCCCC | |
| E3#9 | CCCCCCGACGTTGTGCCCAAGTTGCCCCCC | |
| E3#10 | CCCCCCAAGGGACTGAAAGCAATGCCCCCC | |
| E4#1 | CCCCCCGATCCAACCGCTGCCACACCCCCC | |
| E4#2 | CCCCCCACGACGTACACTCCTCCTCCCCCC | |
| E4#3 | CCCCCCAACCGCATGTACTCTCCGCCCCCC | |
| E4#4 | CCCCCCAACATGCACAGACGTCCGCCCCCC | |
| E4#5 | CCCCCCAACCATGCACAACGCGTGCCCCCC | |
| E4#6 | CCCCCCACACAACCTGCTCCTCCTCCCCCC | |
| E4#7 | CCCCCCCCCCCCCCCCCCCCCCCCCCCCCC | |
| E4#8 | CCCCCCACGCACAATCCGGCACTTCCCCCC | |
| E4#9 | CCCCCCACAGACTGCAGTCATGTGCCCCCC | |
| E4#10 | CCCCCCACACCAGCCACACGTGCGCCCCCC | |
| E5#1 | CCCCCCACGCACCGACAGCACACTCCCCCC | |
| E5#2 | CCCCCCACACCACACCACACCGATCCCCCC | |
| E5#3 | CCCCCCACGACAACCAACACTGTGCCCCCC | |
| E5#4 | CCCCCCAGCACACTACACACGGCGCCCCCC | |
| E5#5 | CCCCCCACACCACCTCACGACGTGCCCCCC | |
| E5#6 | CCCCCCACACCACCAGACACTGCGCCCCCC | |
| E5#7 | CCCCCCACCAACACCAGCCGTGCGCCCCCC | |
| E5#8 | CCCCCCACACACACCACACGTGCTCCCCCC | |
| E5#9 | CCCCCCACACAACACCCGACGCGGCCCCCC | |
| E5#10 | CCCCCCACACACACAACGACGCGGCCCCCC | Same as E6#1 |
| E6#1 | CCCCCCACACACACAACGACGCGGCCCCCC | Same as E5#10 |
| E6#2 | CCCCCCAGCACAACACGGCAACCTCCCCCC | |
| E6#3 | CCCCCCAACACACCACAGACTCTGCCCCCC | |
| E6#4 | CCCCCCACACACCATCAGACGCCGCCCCCC | |
| E6#5 | CCCCCCAGCAGCACACGACACACTCCCCCC | |
| E6#6 | CCCCCCACGCCAACACATTCCGCTCCCCCC | |
| E6#7 | CCCCCCAACACACACAGCCGTCCGCCCCCC | |
| E6#8 | CCCCCCAACACACACAGACGCACGCCCCCC | |
| E6#9 | CCCCCCAGCACCAGACAGCACACTCCCCCC | |
| E6#10 | CCCCCCACCACGATCCTCACTCCGCCCCCC | |

**Table S1**. 10 most frequent ssDNA sequences from round 3, 4, 5, 6 in experimental SELEC groups (E3, E4, E5, E6). #1 to #10 are denoted in the order of descending frequency. Primer region is omitted.

| Name | Sequence |
| --- | --- |
| C3#1 | CCCCCCACAGACCGACGTGTGCTGCCCCCC |
| C3#2 | CCCCCCTGGGAGCCATCTTGTGCGCCCCCC |
| C3#3 | CCCCCCGTTCAGCCTTTTCGTTCGCCCCCC |
| C3#4 | CCCCCCGGAATCTCCGGCGTCTATCCCCCC |
| C3#5 | CCCCCCTAGCACAGGTCGTCTATTCCCCCC |
| C3#6 | CCCCCCGCCAATATAGCCCTTCCGCCCCCC |
| C3#7 | CCCCCCAATCACTGCAATGGTCGTCCCCCC |
| C3#8 | CCCCCCAACACATTGACGTGCACTCCCCCC |
| C3#9 | CCCCCCGGGCTGTGCCGTCATGCGCCCCCC |
| C3#10 | CCCCCCGATGGGGAATCATGCGTGCCCCCC |
| C4#1 | CCCCCCACACAGCATCATTCCGCTCCCCCC |
| C4#2 | CCCCCCGCACCAACCAGCCGTCTGCCCCCC |
| C4#3 | CCCCCCTCACCACATTCGACGGCGCCCCCC |
| C4#4 | CCCCCCACCACAAGTGACTGTCCTCCCCCC |
| C4#5 | CCCCCCGCCGACATGACTCCTCCTCCCCCC |
| C4#6 | CCCCCCACACACCAATGACCTGTGCCCCCC |
| C4#7 | CCCCCCTACCCACACCACACACTGCCCCCC |
| C4#8 | CCCCCCACTGCACATCGACGCGCGCCCCCC |
| C4#9 | CCCCCCATTGCCGCCATCCTCATGCCCCCC |
| C4#10 | CCCCCCAGGCCACCGTCGCACGTGCCCCCC |
| C5#1 | CCCCCCAACACCACACACGGCGCTCCCCCC |
| C5#2 | CCCCCCAGCACACTCCACTCCGCTCCCCCC |
| C5#3 | CCCCCCGCACACACCAGCCGTCTGCCCCCC |
| C5#4 | CCCCCCAACCACACACCGTCCGCTCCCCCC |
| C5#5 | CCCCCCACCACACCATCGACGCGTCCCCCC |
| C5#6 | CCCCCCAGCCACACGACGCGCTCTCCCCCC |
| C5#7 | CCCCCCACGGCACACACCATCGCTCCCCCC |
| C5#8 | CCCCCCACGACACTGCACGACGCGCCCCCC |
| C5#9 | CCCCCCACGGCAACTCCCATTCCGCCCCCC |
| C5#10 | CCCCCCACGACACCACACTGCTCTCCCCCC |
| C6#1 | CCCCCCACCGCATCGACATGTGCTCCCCCC |
| C6#2 | CCCCCCACCGCACGAGCCAGTGTGCCCCCC |
| C6#3 | CCCCCCTCACCACATTCCGCTGTGCCCCCC |
| C6#4 | CCCCCCACCGAGAGCAGACGATGTCCCCCC |
| C6#5 | CCCCCCAACACCACACACGGCGCTCCCCCC |
| C6#6 | CCCCCCGCAGCGTGACTTGACGTGCCCCCC |
| C6#7 | CCCCCCAACACGGCCCTCATGTCGCCCCCC |
| C6#8 | CCCCCCAGCCGTATGCACACCTCACCCCCC |
| C6#9 | CCCCCCACACACCGTTCATCCGCGCCCCCC |
| C6#10 | CCCCCCGCTGATCGACGACACGTGCCCCCC |

**Table S2**. 10 most frequent ssDNA sequences from round 3, 4, 5, 6 in control SELEC groups (C3, C4, C5, C6). #1 to #10 were denoted in the order of descending frequency. Primer region is omitted.

| Name | Sequence |
|---|---|
| pR0 | AGCGTCGAATACCACTACCCCCCCNNNNNNNNNNNNN<br>NNNNNCCCCCCGACCACGAGCTCCATTAG |
| pE6#9 | AGCGTCGAATACCACTACCCCCCCAGCACCAGACAGCAC<br>ACTCCCCCCGACCACGAGCTCCATTAG |
| pC6#8 | AGCGTCGAATACCACTACCCCCCCATTACAGCGGACAAG<br>TGTCCCCCCGACCACGAGCTCCATTAG |
| R0 | CCCCCCNNNNNNNNNNNNNNNNNNCCCCCC |

**Table S3**. ssDNA sequences utilized in experiments: pR0 is the unevolved initial ssDNA library, pE6#9 and pC6#8 are sequences used for investigating the effect of the PCR primer region to the ssDNA-SWNT 5-HT response and solvatochromatic shift assay by sodium cholate, and R0 is the PCR-primer-truncated initial ssDNA library.

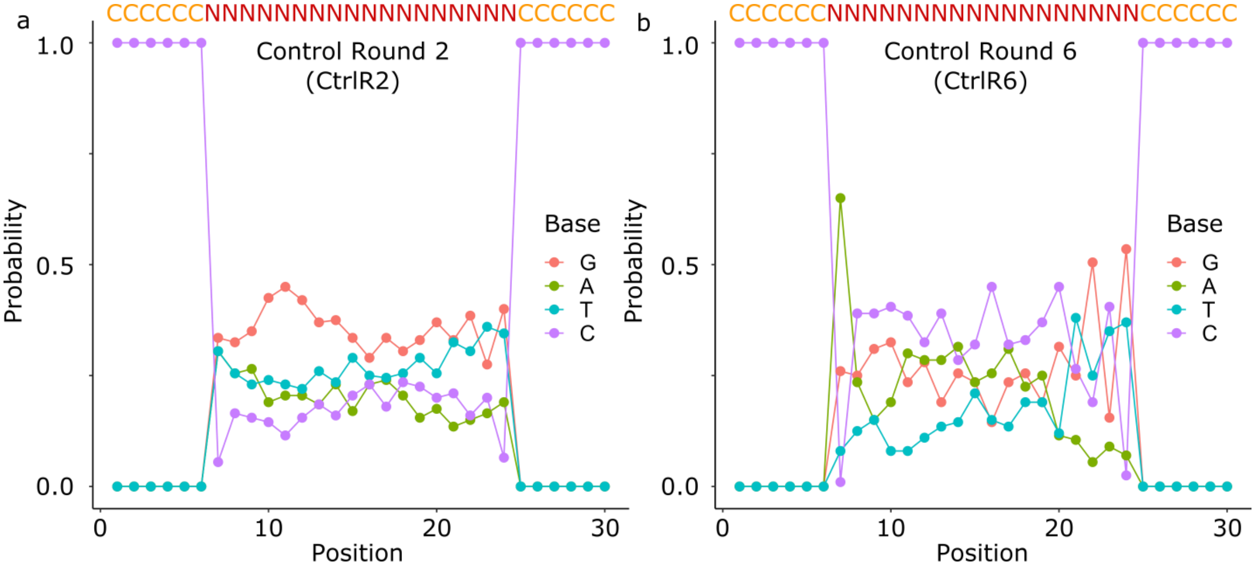


**Figure S1**. **Evolution of nucleotide identity prevalence in the control SELEC library.** Probability of each nucleotide at each position in the control SELEC library calculated from the top 200 sequences in SELEC (a) round 2, and (b) round 6.

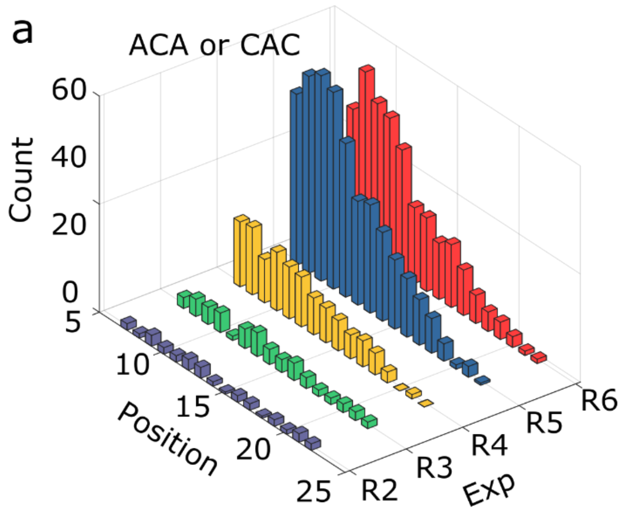


**Figure S2**. **Prevalence of palindromic sequences evolved in SELEC experimental and control libraries.** Palindromic trimer incidences of (ACA)/(CAC) at each nucleotide position for (a) experimental and (b) control SELEC libraries.

**Principal component analysis (PCA) of DNA sequences from SELEC**

Top 200 ranking sequences from round 2,3,4,5, and 6 in experimental and control SELEC libraries were analyzed with PCA. The total number of sequences used for PCA analysis is 2000, and all calculations were performed in Matlab. Nucleotide information of a sequence was converted by using a boolean vector consisting of 0 or 1. For example, ATGC and TACG sequences were digitized as (1000 0100 0010 0001) and (0100 1000 0001 0010), respectively. We utilized only the 18-mer random region of the ssDNA, and did not include the PCR primer nor the $(C)_6$ flanking regions in PCA analysis. PCA was performed by using the Matlab function, pca() with a singular value decomposition (SVD) algorithm. The top 200 ranking sequences from experimental and control SELEC libraries were plotted in terms of principal component scores of the first principal component (PC1) and the second principal component (PC2) (Figure S3).

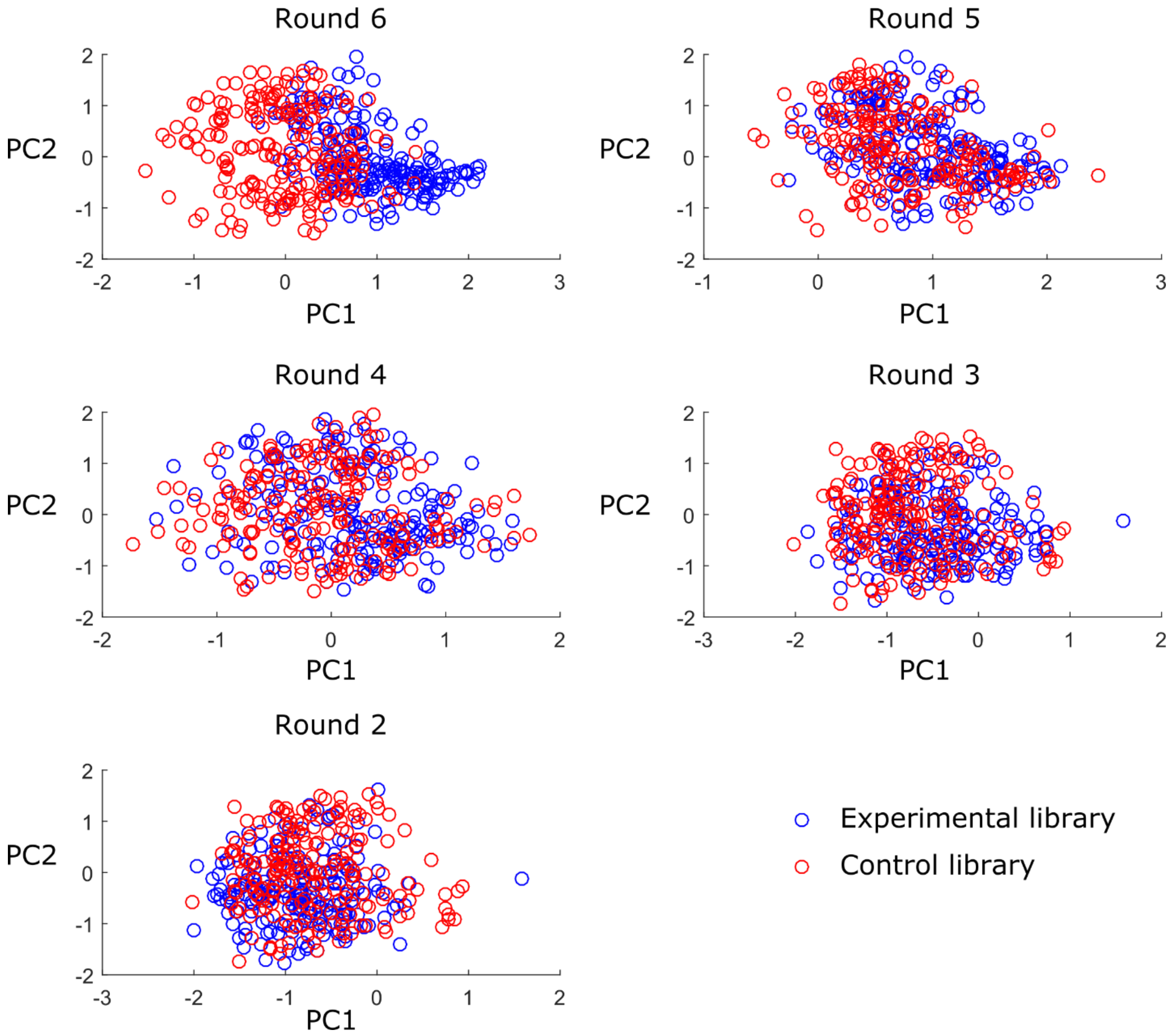


**Figure S3. Scatter plots of principal component 1 (PC1) versus principal component 2 (PC2) for experimental and control SELEC library sequences.** PCA was performed for the top 200 ranked DNA sequences from round 2 to round 6 in the experimental and control SELEC libraries.

**Omission of primer regions from ssDNA-SWNT constructs.**

We hypothesize that the flanking primer regions will have marginal influence on the corona conformation of ssDNAs on SWNT but will putatively reduce the number density of 5-HT recognition sites on SWNT since the 66-mer ssDNA containing the primer domains occupies more space than the primer truncated 30-mer ssDNA. To test this hypothesis, we prepared two ssDNA-SWNT constructs from E6#9 without the primer domain and pE6#9 containing the primer domain, and compared the DNA-SWNT fluorescence response for each upon exposure to 100 µM 5-HT. ssDNA-SWNT constructs revealed distinct peaks in fluorescence spectra, corresponding to different SWNT chiralities[1] (Figure S2). Both DNA-SWNT constructs with and without primer regions showed a fluorescence increase following addition of 100 µM 5-HT, however E6#9-SWNT yielded a fluorescence enhancement of $\Delta F/F_0$ = 1.91, which is twice as high as the $\Delta F/F_0$ = 1.09 of pE6#9-SWNT. We calculate $\Delta F/F_0 = (F - F_0)/F_0$ based on $F_0$ of baseline fluorescence intensity before analyte addition and F of the fluorescence intensity 10 s after 5-HT addition for the (8,6) SWNT chirality (~1195 nm)). Thus, we opted to use ssDNA-SWNT constructs without primer regions for experiments in our manuscript, unless denoted otherwise.

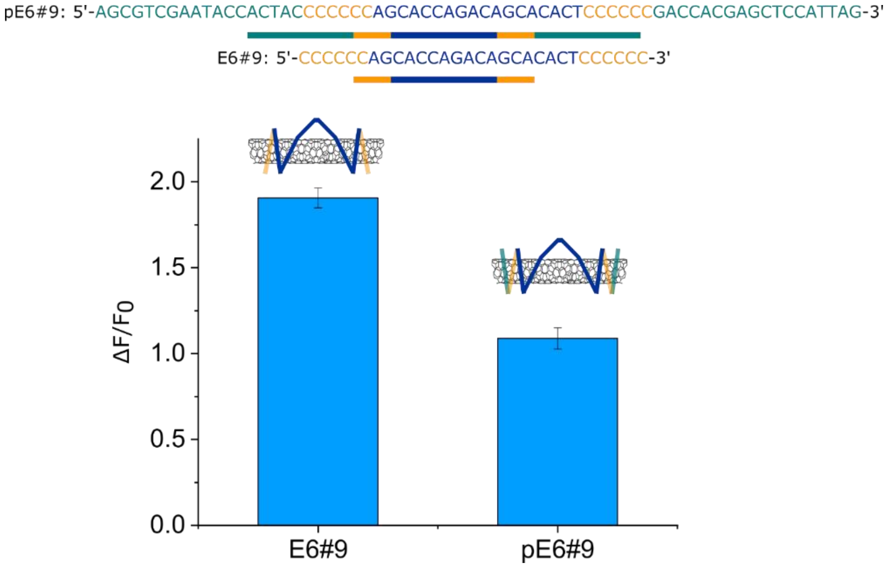


**Figure S4. Truncation of primer region from ssDNA sequence improves the 5-HT response of ssDNA-SWNT.** $\Delta F/F_0$ of pE6#9 DNA-SWNT and primer-truncated E6#9 DNA-SWNT 10 s after addition of 100 µM 5-HT. $\Delta F/F_0$ refers to the FL intensity change at the peak intensity of the (8,6) SWNT chirality (~ 1195 nm). Error bars denote standard deviation from n = 3 trials.

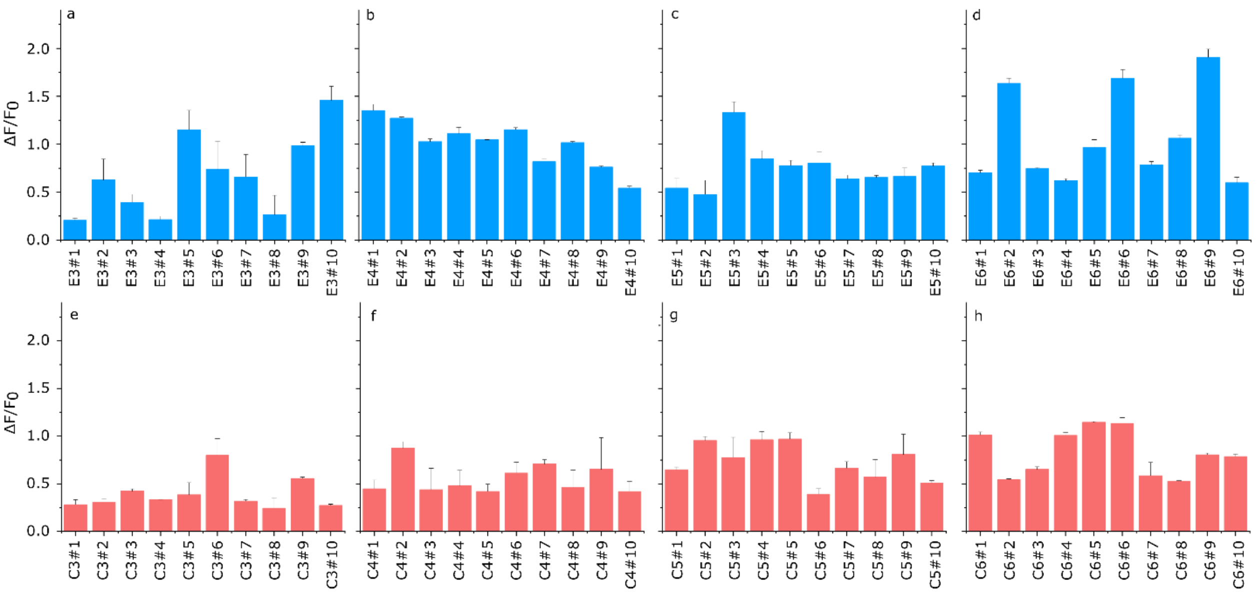


**Figure S5**. **ssDNA-SWNT response to 5-HT from experimental and control SELEC groups.** Top 10 most abundant ssDNA-SWNT sequences from round 3, 4, 5, 6 in experiment (a,b,c,d) and control (e,f,g,h) SELEC groups. $\Delta F/F_0$ refers to the fluorescence intensity change at the peak intensity of the (8,6) SWNT chirality (~1195 nm) 10 s after addition of 100 μM 5-HT. Error bars denote standard deviation from n = 3 trials.

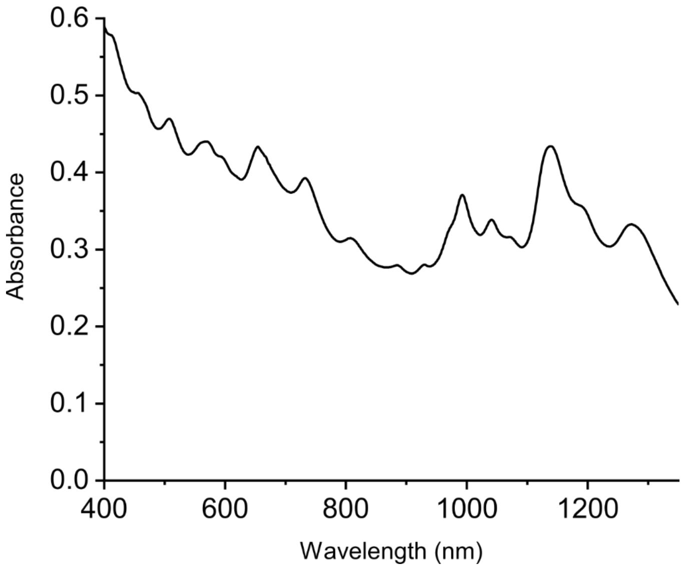


**Figure S6**. **Absorption spectrum of nIRHT in DI water.**

**Time-dependent fluorescence response of nIRHT to 5-HT and other neurotransmitter analytes.**

Time-dependent nIR fluorescence of nIRHT nanosensors following 100 μM 5-HT addition shows a rapid initial fluorescence surge followed by a slow minor fluorescence decay. In contrast, addition of 100 μM dopamine also yields a rapid initial fluorescence surge without a subsequent fluorescence decay (Figure S5 and S6a). A feasible explanation for the fluorescence decay following exposure to 5-HT could be the slow oxidation of 5-HT[2] by SWNT-mediated photoinduced charge transfer[3] since 5-HT is a redox active neurotransmitter. It is known that in ambient condition, 5-HT slowly oxidizes via polymerization among 5-HT monomers, which we confirmed by monitoring the increase in 5-HT absorption at 325 nm over the course of several days[2] (Figure S6b). To test whether we induce irreversible 5-HT oxidation with laser exposure and nIRHT nanosensor co-incubation, we characterized the chemical state of 5-HT following incubation with nIRHT nanosensors and laser exposure commensurate with experimental conditions. First, 10 mg/L nIRHT nanosensors in PBS was prepared and incubated with 100 μM 5-HT under continuous laser excitation at 721 nm. The absorption spectra of nIRHT nanosensor solutions 5-fold diluted in PBS, were measured before and after addition of 5-HT. The absorption profile of 5-HT remained intact and the UV absorption spectra did not reveal the appearance of 5-HT oxidation peak over 60 min of laser excitation (Figure S6c). We next analyzed 5-HT by NMR spectroscopy after nanosensor coincubation to investigate the existence of a potential oxidative side product of 5-HT such as tryptamine dione monomers[4]. Following 30 min of 721 nm laser excitation of 10 mg/L nIRHT nanosensor with 100 μM 5-HT, nIRHT nanosensors were removed by membrane filtration and the flow-through containing the 5-HT and any of its small molecule derivatives was analyzed by NMR. The eluted solution indicated the existence of intact 5-HT and no 5-HT derivatives were observed (Figure S6d, e). These results preclude the presence of irreversible oxidized 5-HT products. The observed slow fluorescence decay kinetics ($\tau_{1/2} > 100$ s) will not prevent the utility of nIRHT imaging 5-HT neuromodulation since 5-HT release and reuptake occurs in the temporal range of milliseconds to seconds[5], and we have shown nIRHT to be reversible (Figure 3).

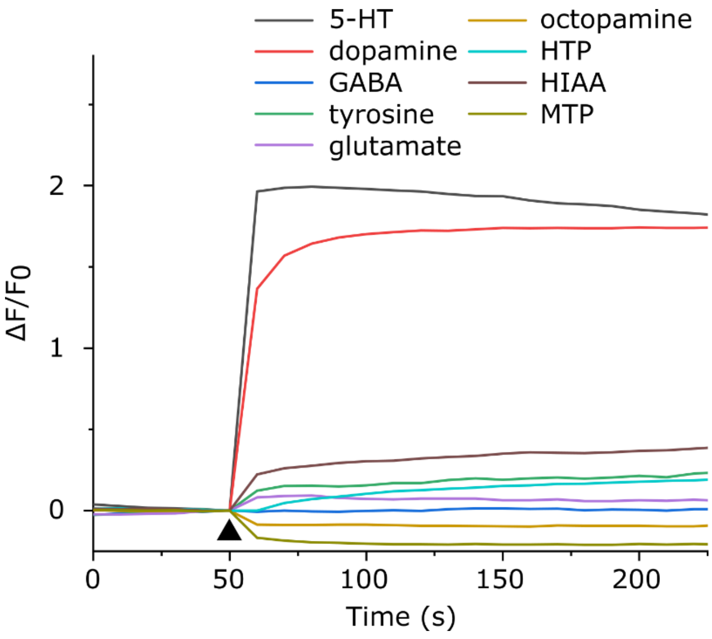


**Figure S7. Time-dependent NIR fluorescence response of nIRHT nanosensor to various neurotransmitter and metabolite molecules.** (a) Representative $\Delta F/F_0$ of nIRHT nanosensor following addition of 100 µM of each analyte at the time indicated by the black triangle. $\Delta F/F_0$ refers to the fluorescence intensity change at the peak intensity of the (8,6) SWNT chirality (~1195 nm). HTP = 5-hydroxy-L-tryptophan, HIAA = 5-hydroxyindoleacetic acid, MTP = 5-methoxytryptamine (MTP), GABA = γ-aminobutylic acid.

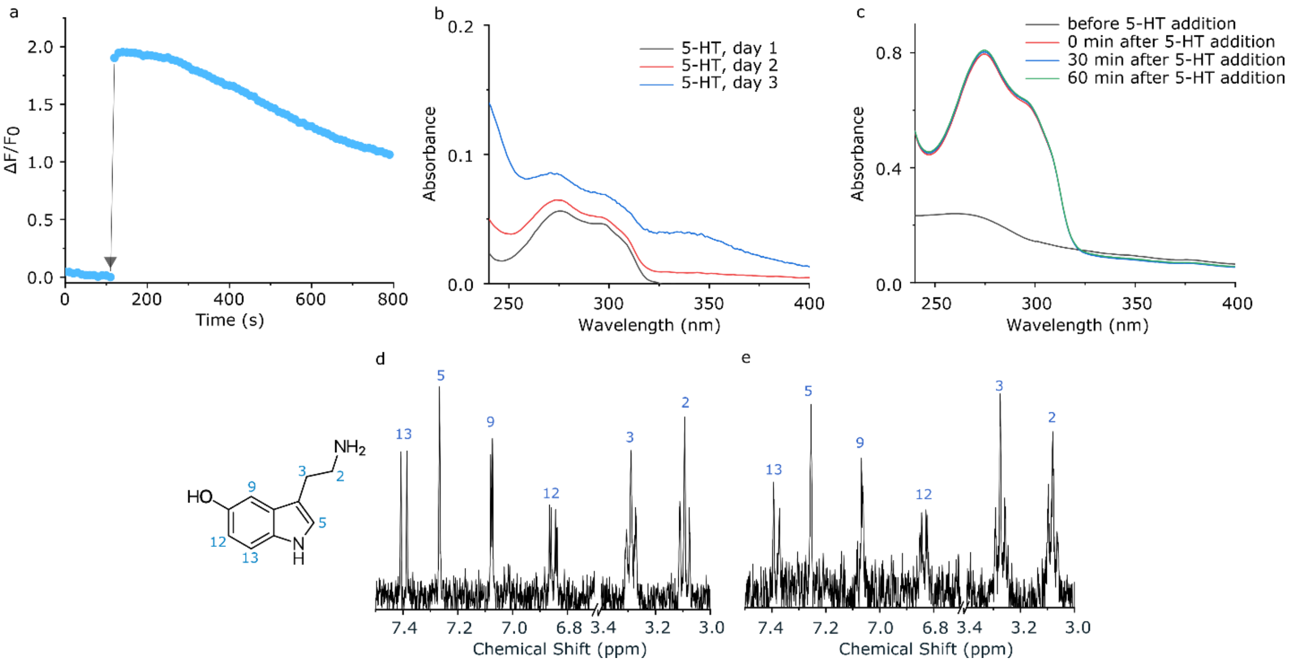


**Figure S8. Fluorescence intensity profile of nIRHT nanosensors following 5-HT addition is not due to 5-HT oxidation.** (a) $\Delta F/F_0$ of nIRHT nanosensor following addition of 100 µM 5-HT at the time indicated by the black triangle. $\Delta F/F_0$ of nIRHT nanosensor shows a rapid fluorescence surge and subsequent slow fluorescence decrease. (b) Absorbance spectra of 10 µM 5-HTsolution in PBS buffer which was stored in ambient conditions and showing the appearance of an absorption band at 325 nm indicating oxidation over the course of 3 days. (c) Absorbance spectra of nIRHT nanosensors before and after addition of 100 µM 5-HT and under continuous exposure to 721 nm laser irradiation, showing no sign of oxidation. 1H NMR (400 MHz, $D_2O$, 298 K) spectra of (d) 100 µM 5-HT and (e) eluted 5-HT after 10 min incubation with nIRHT nanosensor under 721 nm excitation. NMR spectra does not indicate any spectral shift or the appearance of new NMR peaks after co-incubation with nIRHT nanosensor suspension under laser irradiation.

**ssDNA-SWNT stability probed by a solvatochromatic shift assay with sodium cholate-induced surface adsorption**

Our experimental results suggest several ssDNA-SWNT constructs from our SELEC experimental group exhibit 5-HT molecular recognition. Sodium cholate (SC) is a surfactant that is known to induce dynamic surfactant exchange from ssDNA-wrapped SWNT to SC-wrapped SWNT accompanied by a solvatochromic blue shift in the optical transition energy of SWNTs[6, 7]. It has been shown that the kinetics of this solvatochromic shift is inversely proportional to the initial ssDNA-SWNT affinity, and that the ssDNA-SWNT affinity can be enhanced by molecules that exhibit molecular recognition by the ssDNA-SWNT corona[7]. We prepared three ssDNA-SWNT suspensions with the following ssDNAs: the initial random ssDNA library (pR0), the 9th place sequence in the experimental group round 6 which was characterized as our best 5-HT nanosensor (pE6#9), and the 8th place sequence in the control group round 6 (pC6#8). These sequences all contain the PCR primer regions at each end. When incubated with 0.25 wt% SC, all constructs showed time-dependent spectral blue shifts in fluorescence peak wavelengths, which we quantified for the (8,6) chirality at an initial peak value of 1195 nm (Figure S7a). We observed that the pE6#9-SWNT construct exhibited a faster 2 nm spectral shift (58 seconds) than the pR0-SWNT construct (86 seconds) upon addition of 0.25 wt% SC, indicative of a lower stability of the pE6#9-SWNT corona (Figure S7b). Thereafter, we preincubated pE6#9-SWNT and pR0-SWNT constructs with 100 µM 5-HT before adding 0.25 wt% SC for the displacement assay (Figure S7c). Interestingly, for the 5-HT preincubated constructs, pE6#9-SWNT exhibited greater stability as observed by the complete elimination of spectral shift. Conversely, pR0-SWNT constructs preincubated with 5-HT only showed a 94 second spectral shift delay over pR0-SWNT alone, suggesting that 5-HT has higher affinity for pE6#9-SWNT over pR0-SWNT, as expected. The observed 94 second spectral shift delay for pR0-SWNT implies that the initial random ssDNA library also contains some sequences with 5-HT molecular recognition when adsorbed to SWNT.

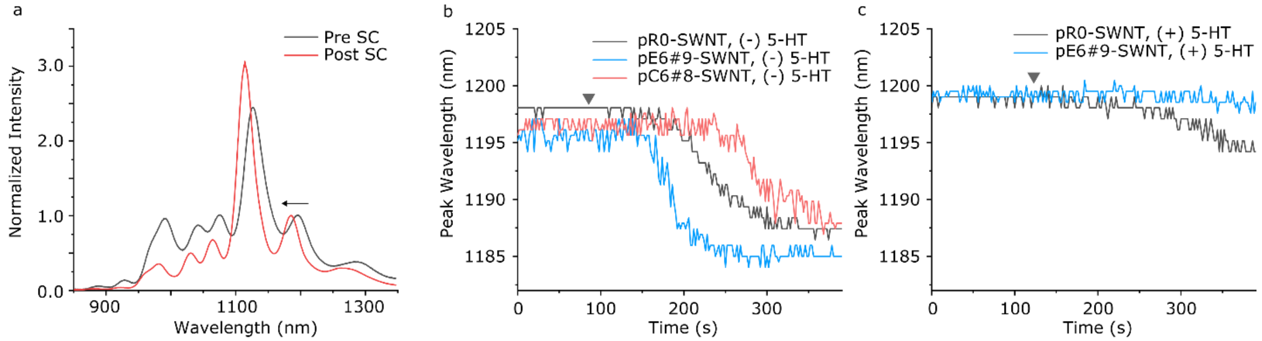


**Figure S9. Solvatochromic spectral shift indicates that SELEC for 5-HT nanosensors selects for ssDNA sequences that have molecular recognition for 5-HT when adsorbed to SWNT.** Three ssDNA-SWNT constructs were prepared in PBS from the initial ssDNA library pool (pR0), the best 5-HT responsive sequence in the SELEC experimental group round 6 (pE6#9), and a similarly-ranked sequence from the SELEC control group round 6 (pC6#8). ssDNA sequences contain both forward and backward primer (p) regions. (a) Normalized fluorescence spectra of pE6#9-SWNT before and after incubation with 0.25 wt% SC in PBS buffer. SC-induced solvatochromic spectral shift is quantified at the (8,6) chirality emission peak (~1195 nm). (b) (8,6) chirality peak wavelength of pR0-, pE6#9-, and pC6#8-SWNT upon addition of 0.25 wt% SC at the time indicated by the gray triangle. (c) (8,6) chirality peak wavelength of pE6#9- and pC6#8-SWNT pre-incubated with 100 μM 5-HT upon addition of 0.25 wt% SC.

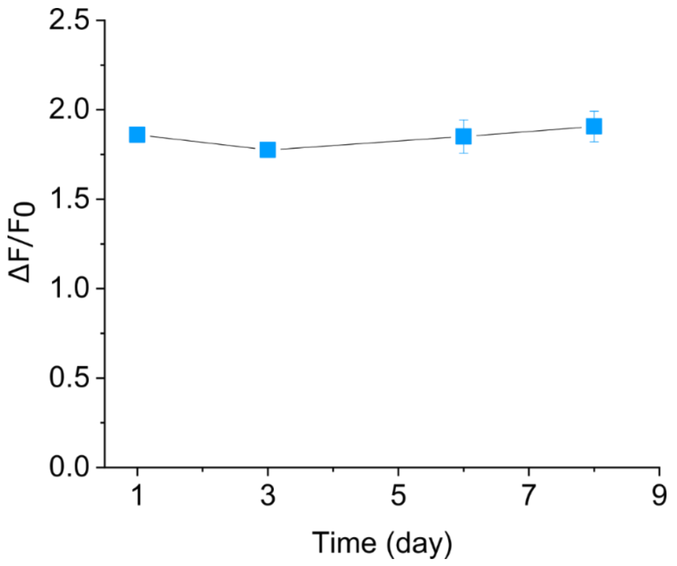


**Figure S10**. **Reproducibility of nIRHT nanosensor fluorescence response to 5-HT.** *$\Delta F/F_0$* of nIRHT nanosensor at 10 s following addition of 100 µM 5-HT measured over a week. nIRHT was stored in PBS buffer at 4 $^{o}$C in the interim. Error bars denote standard deviation from n = 3 trials.

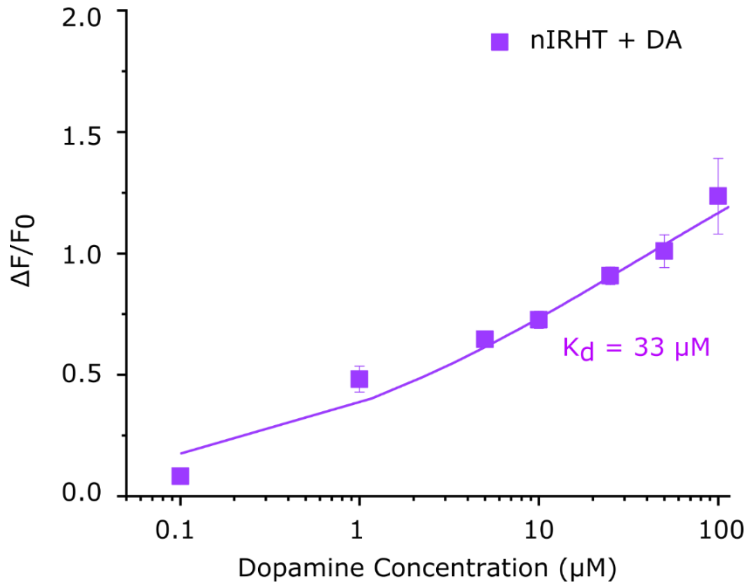


**Figure S11.** ***$\Delta F/F_0$* dopamine concentration dependence of nIRHT.** Error bars are standard deviation from n = 3 independent trials and may be too small to be distinguished in the graph. Experimental data are fitted with the Hill Equation (solid trace).

**Null SWNT response to 5-HT in the absence of ssDNA coating**

We examined the optical response of immobilized bare SWNT without a ssDNA corona phase upon exposure to 10 μM 5-HT. To prepare immobilized bare SWNT, sodium dodecyl sulfate (SDS)-suspended SWNT were prepared and immobilized on an APTES-treated glass slide microfluidic chamber. SDS was removed from the SWNT surface by continuously washing microfluidic chamber with PBS for 1 hour, leveraging the equilibrium between micellular SDS, SDS on the SWNT, and bulk SDS[8]. Addition of 10 μM 5-HT as denoted by a black triangle induced negligible fluorescence modulation of bare SWNTs (Figure S10), confirming that corona phase molecular recognition of ssDNA-SWNTs in this study are a result of ssDNA-induced molecular recognition.

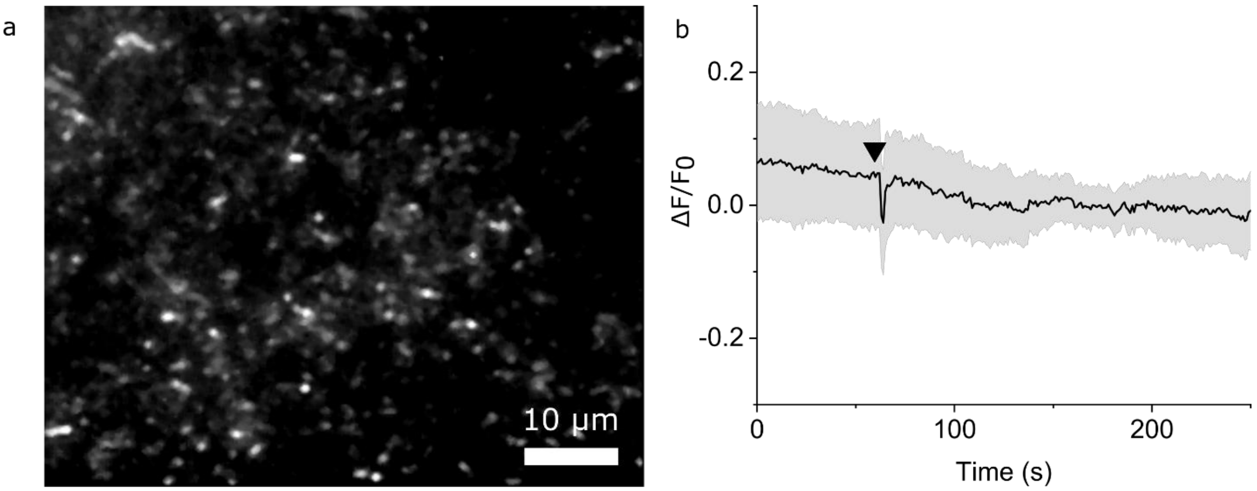


**Figure S12. Null 5-HT fluorescence response of bare SWNT.** (a) NIR fluorescence images of bare SWNT immobilized on the glass surface of a microfluidic microscope slide. (b) *$\Delta F/F_0$* of immobilized SWNTs. Nanosensors were exposed to 10 μM 5-HT (black triangle). Mean traces and standard deviation bands of multiple SWNTs are presented.

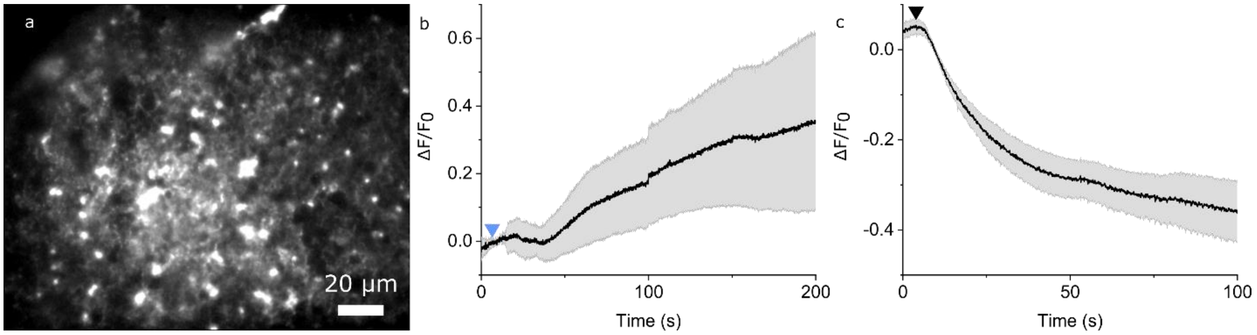


**Figure S13. Time-dependent nIRHT nanosensor response to exogenous 5-HT in acute mouse brain slice.** (a) nIR fluorescence image of nIRHT nanosensors in the dorsal striatum. $\Delta F/F_0$ of nIRHT nanosensors (b) after exogenous addition of 100 µM 5-HT in aCSF (indicated by blue triangle) and (c) after continuous washing with aCSF, initiated at the black triangle. Mean traces and standard deviation bands are of multiple ROIs (n=56). The kinetics in this assay are diffusion-limited by 5-HT introduction into, and clearance from, the imaging chamber.

**nIRHT nanosensor performance in human cerebrospinal fluid**

To assess nIRHT performance in neurologically-relevant conditions, nIRHT was pre-incubated in human cerebrospinal fluid (CSF). A $\Delta F/F_0$ = 0.48 ± 0.02 was obtained following 100 µM 5-HT incubation (Figure S12). The reduced nanosensor response putatively resulted from the blocked 5-HT recognition sites by adsorption of proteins onto the SWNT surface, which form a protein corona[11]. We attribute the attenuated $\Delta F/F_0$ response of nIRHT in slice versus *in vitro* to protein and other biomolecule exposure in acute slice, as suggested by experiments exposing nIRHT to human cerebrospinal fluid (Figure S12). Nonetheless, nanosensor attenuation in brain tissue is a known phenomenon that has not precluded imaging neuromodulatory kinetics in prior work[12-14].

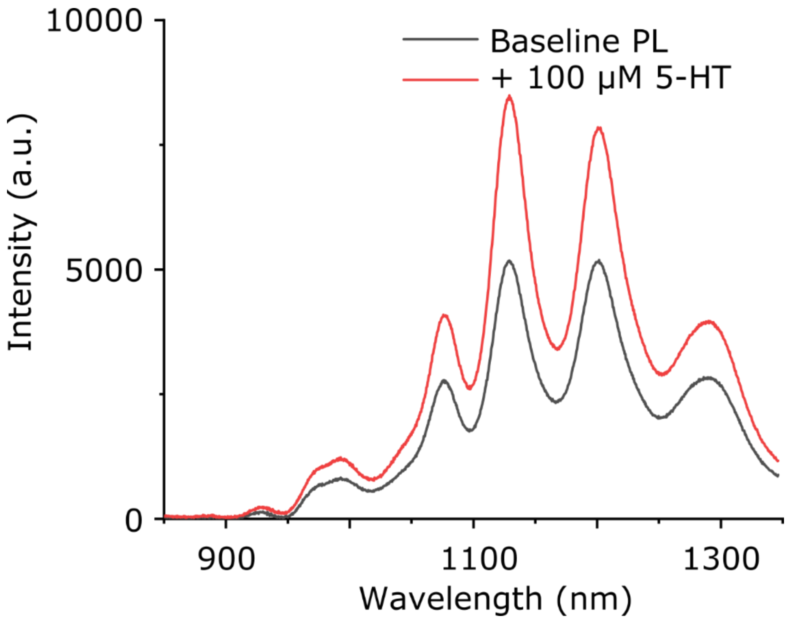


**Figure S14. nIRHT nanosensor response when pre-incubated in cerebrospinal fluid (CSF).** Fluorescence spectra of nIRHT, pre-incubated in CSF for 30 min, excited at 721 nm before (black trace) and 10 s following addition of 100 µM 5-HT (red trace).

**Supplementary Movie Captions**

**Supplementary Movie 1. Time-dependent 5-HT-induced response of nIRHT in microfluidic chamber.** Near-infrared imaging of individual surface-immobilized nIRHT showed reversible fluorescence modulation upon 5-HT addition. Frame rate = 20 frames/s.

**Supplementary Movie 2. nIRHT-labeled brain dorsomedial striatum slice without external stimuli.** Near-infrared imaging of nIRHT-labeled acute mouse striatal brain slice in aCSF. Fluorescence modulation relative to the time-zero image ($\Delta F/F_0$) is shown. Color scale corresponds to the scale bar in Figure 3i,j. Frame rate = 20 frames/s.

**Supplementary Movie 3. nIRHT-labeled brain dorsomedial striatum slice after following addition of 100 μM 5-HT, and subsequent washing by aCSF.** Near-infrared imaging of nIRHT-labeled acute mouse brain slice showed a fluorescence surge following bath application of 100 μM 5-HT at ~10 s. The brain slice was subsequently washed by continuous aCSF flow (washing start indicated by a separate time stamp that re-starts at 0 s). aCSF washing induced nIRHT fluorescence return to baseline. Image analysis and video settings are the same as for Supplementary Movie 2.
